# Double knockout of rice *OsVIT1* and *OsVIT2* genes reveals a trade-off between iron biofortification and iron excess tolerance

**DOI:** 10.1101/2025.06.04.657710

**Authors:** Betina Debastiani Benato, Angie Geraldine Sierra Rativa, Raquel Vargas Olsson, Yugo Lima-Melo, Eduardo Santos, Gabriel Sgarbiero Montanha, Jover da Silva Alves, Lucas Roani Ponte, Victor Hugo Rolla Fiorentini, Fernando Mateus Michelon Betin, Francieli Ortolan, João Paulo Rodrigues Marques, Raul Antonio Sperotto, Hudson Wallace Pereira de Carvalho, Stefano Cesco, Tanja Mimmo, Raphael Tiziani, Gian Maria Beone, Nathalia Navarro, Hannetz Roschzttardtz, Carlos Alberto Pérez, Ricardo Fabiano Hettwer Giehl, Marcia Margis-Pinheiro, Felipe dos Santos Maraschin, Felipe Klein Ricachenevsky

## Abstract

Rice (*Oryza sativa* L.) is a staple food for half of the world’s population, but lacks essential nutrients such as iron (Fe). Fe deficiency is one of the most common nutritional problems in humans, and biofortification of rice grains is a cost-effective approach to deliver more Fe to people’s diet. Two Vacuolar Iron Transporters, OsVIT1 and OsVIT2, were shown to negatively regulate Fe translocation to rice developing panicles, as single mutants *osvit1* and *osvit2* have increased Fe concentration in seeds. Importantly, rice plants are frequently cultivated in waterlogged soils that are highly reductive and prone to Fe^3+^ reduction to the more soluble Fe^2+^, which can accumulate and cause Fe toxicity. Little is known about which genes control Fe excess detoxification. OsVIT1 and OsVIT2 transport Fe into the vacuole, and *OsVIT2* is induced under Fe excess, but whether they play a role in Fe detoxification was not demonstrated. We generated double mutants *osvit1osvit2* using CRISPR-Cas9 to test whether loss of function of both genes could increase Fe concentration in seeds, and to test whether their loss of function has impact in rice Fe excess tolerance. We showed that *osvit1osvit2* double mutants accumulated more Fe in brown rice. Fe accumulation was clear in embryo scutellum and plumule, suggesting VIT transporters have a role in determining Fe spatial distribution. We also showed that root uptake contributed significantly for increased Fe accumulation in *osvit1osvit2* seeds, suggesting OsVIT1 and OsVIT2 are involved in sequestering Fe in vegetative tissues and decreasing translocation. Strikingly, we found that *osvit1osvit2* plants were more sensitive to Fe excess, revealing a trade-off between Fe biofortification and Fe excess tolerance. Our data indicates OsVIT1 and OsVIT2 are key for Fe excess detoxification, which should be considered in their use as targets for biofortification.

## Introduction

Rice (*Oryza sativa* L.) is a key crop for maintaining global food security, as it is a staple food for nearly half of the world’s population and provides around 20% of all calories consumed by humans (Elert, 2014; Fukagawa & Ziska, 2019). Despite its importance as a calorie source, rice is poor in essential micronutrients, such as iron (Fe) and zinc (Zn), making people who rely on it prone to malnutrition. This combination of energy-dense food with inadequate nutrient intake is known as “hidden hunger”, which is estimated to affect at least two billion people worldwide (Allen, 2006; Gödecke *et al*., 2018). Fe and Zn are crucial for human health, and their lack is among the most common dietary deficiencies, affecting mainly children and women in low-income, developing countries, leading to several health-related issues (Stevens *et al*., 2022). While cereals generally have low concentrations of these micronutrients, rice has the lowest baseline and the least genetic variability for these traits compared to other cereals (Garcia-Oliveira *et al*., 2018; Wairich *et al*., 2022), making conventional breeding challenging. Therefore, genetic engineering is the most promising approach for rice biofortification, aiming to enhance the concentrations of these nutrients in the edible parts at concentrations that can impact human nutrition (Wairich *et al*., 2022).

Fe is also essential for plants, as it plays a key role in electron transport in both chloroplast and mitochondria due to its ability to switch from Fe^2+^ to Fe^3+^ oxidation states. It also participates in chlorophyll and DNA synthesis. Despite its abundance in soils, Fe is often not readily available for root uptake (Colombo *et al*., 2014). To ensure sufficient Fe supply and avoid deficiency, plants have evolved distinct mechanisms to solubilize Fe in the soil, take it up by root transporters, and to maintain adequate distribution within organs, tissues, and cells (Chao & Chao, 2022; Rodrigues *et al*., 2023; Spielmann *et al*., 2023; Huang *et al*., 2024). These mechanisms are classically separated into Strategy I, or reduction-based, found in all non-Poaceae species; and Strategy II, or chelation-based, common to the Poaceae family (Kobayashi & Nishizawa, 2012; Chao & Chao, 2022). More recently, coumarin secretion to the rhizosphere, described in *Arabidopsis thaliana* but also observed in other eudicot species, has been shown to be important for Fe solubilization and reduction (Tomasi *et al*., 2008; Rajniak *et al*., 2018; Robe *et al*., 2021; Paffrath *et al*., 2024). Rice and its closely related wild relatives employ a combined strategy, consisting of a full Strategy II plus the ability to transport Fe^2+^ (Ishimaru *et al*., 2006; Wairich *et al*., 2019). Thus, plants use complex mechanisms to tightly maintain Fe homeostasis and avoid Fe deficiency.

Fe can become toxic when accumulated at high concentrations, leading to oxidative stress due to Fenton chemistry, causing tissue damage (Wairich *et al*., 2024). Fe toxicity is particularly common in rice plants, as they are often cultivated in waterlogged soils. Under these conditions, the soil solution becomes more reductive and causes Fe^3+^ change to the more soluble Fe^2+^, which is transported into root cells and can accumulate to harmful levels (Aung & Masuda, 2020). Fe toxicity can reduce rice yields by 12% to 100%, depending on the cultivar and edaphic conditions (Sahrawat, 2005). In contrast to Fe deficiency, Fe excess is less well understood mechanistically, and only a few genes have been linked to Fe excess detoxification or tolerance mechanisms in rice (Stein *et al*., 2009; Wu *et al*., 2019; Aung *et al*., 2019; Li *et al*., 2019). Therefore, there is a need to improve our understanding of how rice plants cope with Fe excess.

Members of the Vacuolar Iron Transporter (VIT) family are promising candidate genes for both biofortification and Fe excess detoxification. The first gene described in this family, *AtVIT1* from *Arabidopsis thaliana*, encodes a protein critical to maintain the distribution of Fe in seeds (Kim *et al*., 2006). Rice has two *AtVIT1* homologs, *OsVIT1* and *OsVIT2*. Both are Fe transporters localized to the tonoplast, and are involved in Fe delivery to seeds (Zhang *et al*., 2012). Single *osvit1* and *osvit2* mutants showed increased Fe concentration in brown rice and embryos, while Fe levels were also elevated in phloem sap but lower in flag leaves compared to wild type (WT) (Zhang *et al*., 2012). These findings suggest that both transporters sequester Fe in flag leaf vacuoles, reducing Fe availability for translocation to developing seeds. Interestingly, Fe spatial distribution was disrupted in embryos of both single mutants (Zhang *et al*., 2012; Bashir *et al*., 2013). More recent work showed that OsVIT2 is involved in sequestering Fe in the mestome sheath, nodes, and aleurone layer of seeds. A CRISPR/Cas9-generated *osvit2* loss-of-function mutant had increased Fe concentration in the endosperm but decreased Fe in the aleurone layer, without affecting yield (Che *et al*., 2021). Additionally, an *osvit2* generated by ethyl methyl sulfonate (EMS) mutagenesis showed increased Fe levels in seeds (Kandwal *et al*., 2022). Thus, mutations in either *OsVIT1* or *OsVIT2* are strong candidates for biofortification, as both increase Fe concentration in target tissues. Moreover, phenotypes observed in single mutants as well as gene expression patterns suggest that *OsVIT1* and *OsVIT2* are not redundant. *OsVIT2* is consistently up-regulated in roots of rice plants exposed to Fe excess, whereas *OsVIT1* does not (Zhang *et al*., 2012; Bashir *et al*., 2013; Che *et al*., 2021; Wairich *et al*., 2024). Therefore, considering a possible role of detoxification involving Fe vacuolar sequestration indicates that mutants for OsVIT1 and OsVIT2 could be less tolerant to Fe excess.

Here we generated CRISPR-Cas9 double *osvit1osvit2* knockout mutants to test whether they have increased Fe concentration in seeds and how they tolerate Fe excess. We found that *osvit1osvit2* mutants have higher Fe concentration in seeds, with increased Fe abundance in the and altered distribution in the embryo, particularly in the scutellum and plumule. However, *osvit1osvit2* mutants show reduced tolerance to Fe excess, demonstrating that VIT transporters play a role in Fe excess detoxification. Therefore, our data reveal a trade-off when OsVIT1 and OsVIT2 are mutated, with increased Fe concentration in seeds but decreased tolerance to Fe excess.

## Materials and Methods

### Generation of *osvit1osvit2* plants

An expression cassette (*Hind*III/*Xho*I fragment) containing the sgRNA scaffold and the SpCas9 coding region from *pRGE30* (Xie & Yang, 2013) was subcloned into the *Hind*III/*Sal*I sites of the binary vector *pH7WG2* (Karimi *et al*., 2002). The resulting plasmid was named pH7WG2-Cas9U3pgRNA. We designed a single gRNA targeting both *OsVIT1* and *OsVIT2* genes. Genomic sequences of *OsVIT1* (LOC_Os04g38940) and *OsVIT2* (LOC_Os09g23300) were aligned using ClustalW, and regions with high similarity were selected for sgRNA design using CRISPOR (http://crispor.tefor.net/). The target site CCATCTCCATGGGCCTCGGA was selected as a gRNA directed to the end of the first exon in both genes (Fig 1). Double-stranded 5’-phosphorylated oligos (Table S1) were ligated into pH7WG2-Cas9U3pgRNA (linearized by BsaI). The resulting construct was transformed into rice (*Oryza sativa* ssp. *japonica* cv. Nipponbare) calli by *Agrobacterium*-mediated transformation (Ozawa, 2009), with minor modifications (Ortolan *et al*., 2021).

**Figure 1:**
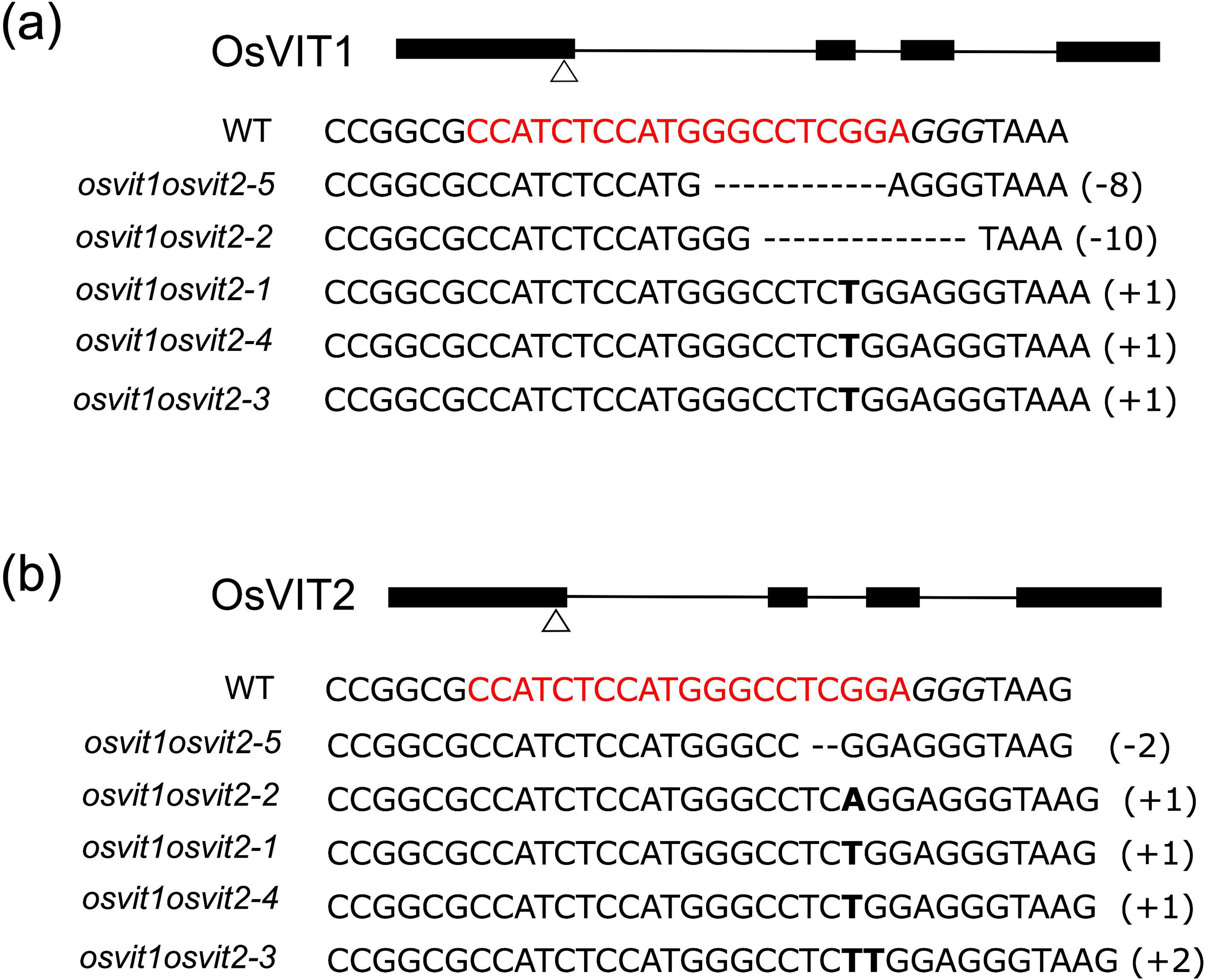
CRISPR-Cas9-mediated editing of *OsVIT1* and *OsVIT2*. *OsVIT1* and *OsVIT2* gene structures (black blocks - exons; black lines - introns). Arrowheads show the gRNA target site at the end of exon I. The WT DNA gene sequence highlights the gRNAs target site (red) and protospacer adjacent motif (PAM; italics). The gene sequences of the five mutant lines are shown. Dashes indicate deletions, and bold letters indicate insertions. Numbers (+/-) indicate the number of nucleotides involved.

Regenerated T1 plants were genotyped by PCR amplicon sequencing. Genomic DNA was extracted from leaves of 10 transformation events and PCR amplifications were carried out using the oligos listed in Supplemental Table 1. PCR products of 230 and 150 bp, respectively, were sequenced and analyzed using TIDE (Tracking of Indels by Decomposition (Brinkman *et al*., 2014). T-DNA-free segregant T2 plants were selected from five independent double homozygous *osvit1osvit2* mutants, named *osvit1osvit2-1* to *osvit1osvit2-5*, to carry on the phenotypic and ionomic analyses.

### Plant Materials and Treatments

Seeds of rice (*Oryza sativa* ssp. *japonica*) cv. Nipponbare (hereafter WT – wild type) and *osvit1osvit2* mutant lines were surface sterilized and germinated in Petri dishes containing filter paper soaked in distilled water. After five to seven days, seedlings were transferred to vermiculite for ∼2 weeks. At the V3 stage (Counce *et al*., 2000), plants were transferred to 400 mL plastic cups with plant holders. Cups were filled with nutrient solution as described previously (Ricachenevsky *et al*., 2011). The control nutrient solution contained 700 μM K_2_SO_4_, 100 μM KCl, 100 μM KH_2_PO_4_, 2 mM Ca(NO_3_)_2_, 500 μM MgSO_4_, 10 μM H_3_BO_3_, 0.5 μM MnSO_4_, 0.5 μM ZnSO_4_, 0.2 μM CuSO_4_, 0.01 μM (NH_4_)_6_Mo_7_O_24_, and 100 μM Fe^+3^-EDTA, pH 5.4 without aeration. Plants were grown at 25 °C ± 1 °C and 180 µmol photons m^-2^ s^-1^ under 16h/8h light/dark photoperiod. The nutrient solution for Fe excess treatments was supplemented with FeSO_4_ (1 mM or 5 mM), while Fe was omitted in the Fe deficiency treatment. Nutrient solutions were replaced every two days.

### Growth and Agronomic Traits

For agronomic traits analyses, plants from WT and *osvit1osvit2* mutant lines were cultivated under greenhouse conditions until maturity. Plant height was measured with a ruler (n = 11-30). The thousand seed weight (here referring to the caryopsis, which is the seed after dehusking the palea and lemma) and plant dry weight were measured using a precision scale (n = 7). The number of spikelets and filled/empty seeds were manually counted (n = 3-7).

### Gas exchange analysis

The CO_2_ assimilation rate (A) and stomatal conductance (*g_s_*) were measured between the fifth and tenth hour after the start of the 16-h light photoperiod using a portable infrared gas analyzer (LCpro-SD, ADC BioScientific, Hoddesdon, UK). Measurements were conducted on the youngest fully expanded leaf, with each sample being measured for 8 minutes (n = 4). Internal conditions within the IRGA chamber during gas exchange measurements were maintained at 1,000 µmol m^-2^ s^-1^ photosynthetic photon flux density (PPFD), 420 ppm CO_2_, 27°C, and approximately 80% relative ambient humidity.

### ICP-MS analyses

Leaf, root and seed samples were collected, rinsed twice in milli-Q water, and dried at 80°C. ICP-MS analyses were performed using two different methods in independent experiments. Seeds, roots, or leaves were weighed into PTFE tubes and digested in HNO_3_ under pressure using a microwave digester (UltraCLAVE IV; MLS). Mineral elemental analysis of digested samples was performed by sector field high-resolution ICP-MS (ELEMENT 2; Thermo Fisher Scientific). Element standards were prepared from certified reference materials from CPI international. For seeds in the Fe supply experiment, the method used was the previously described (Lahner *et al*., 2003) with few adaptations (Huang *et al*., 2019).

### 57Fe experiments

Plants of WT, *osvit1osvit2-1* and *osvit1osvit2-2* were grown in hydroponic nutrient solution under a photoperiod of 11h/13h light/dark (24,000 lux, LED) in Bolzano, Italy. When plants reached anthesis, two sets of treatments were performed using biological quadruplicates. In the first experiment, ^57^Fe root uptake was tested by transferring plants to a 50 mL nutrient solution containing 10 µM of ^57^Fe^3+^-EDTA, prepared from ^57^FeCl_3_ (ISC Science, Oviedo, Spain), until the solution was fully absorbed. The solution was then replaced with the regular nutrient solution until seed set and sampling. In the second experiment, plants grown in the regular nutrient solution were used for flag leaf ^57^Fe supplementation. This was achieved by immersing the flag leaf tip in a microtube containing 1 mL of either 10 µM or 100 µM ^57^Fe^3+^-EDTA nutrient solution, prepared as described above. The seeds were harvested for ^57^Fe quantification as described (Valentinuzzi *et al*., 2020).

### Perls and Perls/DAB staining

Rice seeds from soil-grown plants (WT and *osvit1osvit2* mutant lines) were stained with a freshly prepared 4% (v/v) HCl and 4% (w/v) potassium ferrocyanide solution, as described by Zhang et al. (2012). Perls/DAB staining was carried out as described (Roschzttardtz *et al*., 2009; Vargas & Roschzttardtz, 2023). The seeds were immersed in the staining solution and vacuum-infiltrated for approximately 45 minutes, then longitudinally cut into two halves with a sharp razor blade. Imaging was performed using a Leica M165 FC stereomicroscope, with LAS software.

### X-ray fluorescence spectroscopy (XRF) measurements

Benchtop XRF analyses: the elemental composition and distribution of rice seeds was investigated using microprobe X-ray fluorescence spectroscopy (µ-XRF). Cross-sections of seeds of wild-type (WT) and mutant (*osvit1osvit2-1* and *osvit1osvit2-2*) plants were irradiated by a 30-µm polycapillary optics-focused X-ray beam generated by a Rh anode operating at 45 kV and 900 µA, with an Al 250 µm-thick primary filter selected. The median region of the seeds was scanned using a *ca*. 2 µm length 32-point linescans. Furthermore, at least three points randomly selected at either the aleurone layer or the endosperm tissues were also investigated. In all cases, the spectra were recorded for 10 s by a silicon drift detector (SSD), with a dead-time smaller than 13%. All the analyses were carried out using at least three (n = 3) independent replicates, and only the XRF characteristic intensities above the instrumental limit of detection (ILOD), calculated according to Eq. 1 described below, were considered valid. BG (counts s^-1^) is the elemental background counting rate and t (s) is the acquisition time.

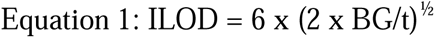

Synchrotron-based XRF analyses at the Canadian Light Source: rice seeds were embedded in Optimal Cutting Temperature compound (OCT) and water, then thinned on both sides to maintain a central section of approximately 100 µm thickness. The seed sections were placed on a solid OCT surface and covered with water and immediately frozen using liquid nitrogen, creating a block with two distinct phases. The water phase of the block was trimmed to the middle of the seeds, and a new layer of water was applied to the trimmed surface. After trimming both sides, the slices were allowed to defrost at room temperature to facilitate their extraction. Finally, the samples were placed on synchrotron tape and covered with a polypropylene film of 5 µm thickness. Samples were fixed in the beamline sample holder using Kapton tape.

The analysis was performed at the BioXAS beamline of the Canadian Light Source (CLS, Saskatoon, Canada) in micro-mode. The beam was focused to 5 (H) x 5 (V) using Kirkpatrick-Baez (KB) mirrors. The beamline was configured for fluorescence mode, with the sample stage positioned at a 45° angle relative to the detector and X-ray beam. A chemical map of the germ, bran, and part of the endosperm of rice seeds was obtained using incident energy of 10 keV, with an acquisition time of 50 ms per pixel, 10 µm step size, and a pixel size of 10 µm. The XRF maps were normalized using the signal recorded by the I ion chamber (positioned before the samples) in PyMca software (Solé *et al*., 2007). Subsequently, the data were further normalized by absorbance using OriginLab (version 2022b, USA), utilizing signals from both the ion chamber before (I) and the ion chamber after (I) the sample in PyMca.

Synchrotron-based XRF analyses at the Brazilian Synchrotron Light Laboratory: rice seeds from WT and mutants (*osvit1osvit2-2*) were first placed under a stereomicroscope and held with the aid of flat tweezers to ensure stability of the sample. Using a sharp steel blade, seeds were trimmed at both sides, then sectioned longitudinally at the middle of the seed from the apical end to the basal end to expose the embryo. The longitudinal median sections of the rice seed with *ca*. 150 µm were fixed into proper XRF sample holders using a Kapton tape, covered with a 5-µm thick polypropylene thin-film and transferred to the Tarumã endstation of the Carnauba - Coherent X-ray Nanoprobe Beamline (Tolentino *et al*., 2023) of the Brazilian Synchrotron Light Laboratory (SIRIUS) at Campinas, Brazil. The samples were irradiated by a 150-nm monochromatic X-ray beam provided by a Kirkpatrick−Baez (KB) achromatic optics at a 9750 eV excitation energy. The XRF mappings were obtained in flyscan mode either at 5 µm or 500 nm lateral resolutions, and the spectra were recorded by two four-element silicon drift detectors (SDD, Vortex-ME4, Hitachi High-Technologies Science America, USA). All the analyses were carried out at room temperature. The elemental intensities detected in each pixel unit were normalized by the ring current (I). The data were then fitted and processed using the PyMCA software (Solé *et al*., 2007).

### Statistical analyses

Statistical analyses were performed as previously described (Ponte *et al*., 2025). Briefly, data normality and homoscedasticity were assessed using the Shapiro-Wilk and Levene’s tests, respectively. When both assumptions were met, comparisons between mutants and WT were performed using Student’s *t*-test. If homoscedasticity was violated, Welch’s *t*-test was used. When data were not normally distributed, the non-parametric Mann-Whitney *U* test was applied. A *P*-value ≤ 0.05 was considered statistically significant.

## Results

### Simultaneous knockout of *OsVIT1* and *OsVIT2* genes

CRISPR/Cas9-mediated targeted mutagenesis was used to generate five independent double knockout mutants named *osvit1osvit2-1* to *osvit1osvit2-5*, using a single gRNA targeting both *OsVIT1* and *OsVIT2* genes. A highly conserved sequence at the end of the first exon was selected as the target site (Fig 1). Sequencing of the target site in T3 plants confirmed that all mutations resulted in truncated protein versions, leading to complete loss of function of both genes (Fig 1; Fig S1). Thus, all five lines are double knockouts for the two rice *VIT* genes.

### *osvit1osvit2* plants are more sensitive to Fe excess

Vacuolar transporters are frequently involved in metal detoxification (Ricachenevsky *et al*., 2018). Although OsVIT1 and OsVIT2 have been identified as vacuolar Fe transporters, their role in excessive Fe sequestration in rice has not been demonstrated. Evidence indicates that *OsVIT2* is consistently induced under Fe excess conditions (Stein *et al*., 2019; Wairich *et al*., 2021, 2024). To test whether rice VIT transporters have a role in Fe excess tolerance, we exposed WT and *osvit1osvit2* mutant plants to high Fe concentration. Leaves from WT and mutant plants were similar in control conditions. However, after three days of growth under 5 mM Fe, *osvit1osvit2* leaves already exhibited clear bronzing spots, characteristic of Fe excess, which were absent in WT leaves (Fig 2a and 2b). After seven days of treatments, *osvit1osvit2* leaves showed severe bronzing spots with necrosis and chlorosis, whereas WT leaves started to show Fe excess symptoms (Fig 2c and 2d). This phenotype was uniform in all five *osvit1osvit2* lines (Fig S2). Therefore, we concluded that *osvit1osvit2* plants were more sensitive to Fe excess compared to WT.

**Figure 2.**
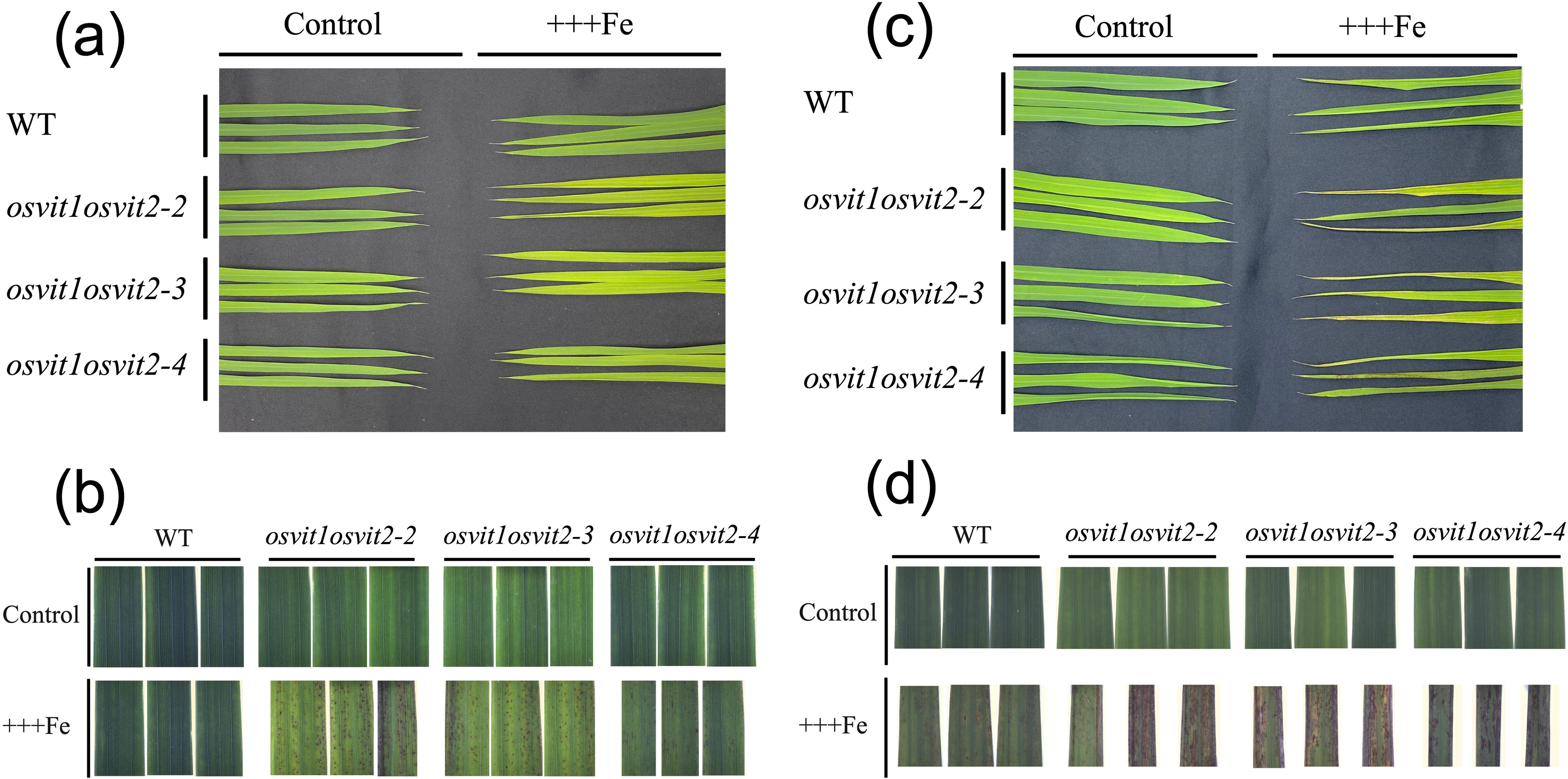
*osvit1osvit2* double knockout mutants are more sensitive to iron excess. (A) Leaves of wild type (WT), *osvit1osvit2-2*, *osvit1osvit2-3* and *osvit1osvit2-4* plants under control (100 µM Fe) and +++Fe (5mM Fe excess) after three days of treatment. (B) Detail showing bronzing spots of leaves shown in (A). (C) Leaves of wild type (WT), *osvit1osvit2-2*, *osvit1osvit2-3* and *osvit1osvit2-4* plants under control and 5m Fe excess after seven days of treatment. (D) Detail showing bronzing spots of leaves shown in (C).

We also observed slight differences in culm height (Fig S3a and S3b), shoot dry weight (Fig S3c and S3d), root length (Fig S3e and S3f), root dry weight (Fig S3g and S3h) and SPAD values (Fig S3i and S3j) comparing the genotypes under control or Fe excess after seven days of treatment. These differences tended to be intrinsic to two out of three lines, and were not consistently affected by Fe excess. Therefore, short-term Fe excess does not significantly affect growth of *osvit1osvit2* plants.

### Photosynthesis is affected in *osvit1osvit2* mutant plants exposed to Fe excess

We tested whether the increased sensitivity of *osvit1osvit2* lines to Fe excess was accompanied by an impact on photosynthesis. As expected, after three days of Fe excess treatment, the net photosynthesis recorded from WT and *osvit1osvit2* mutant plants was lower than under control conditions (Fig 3a and 3b). When comparing WT and mutant plants, we found no difference in control conditions (Fig 3c). However, under Fe excess, *osvit1osvit2* plants had markedly decreased net photosynthesis (Fig 3d). Similar trends were observed for stomatal conductance, with *osvit1osvit2* genotypes exhibiting lower values compared to WT (Fig 3e and 3f). Therefore, photosynthesis and stomatal conductance of *osvit1osvit2* lines are more significantly affected by Fe excess compared to WT.

**Figure 3.**
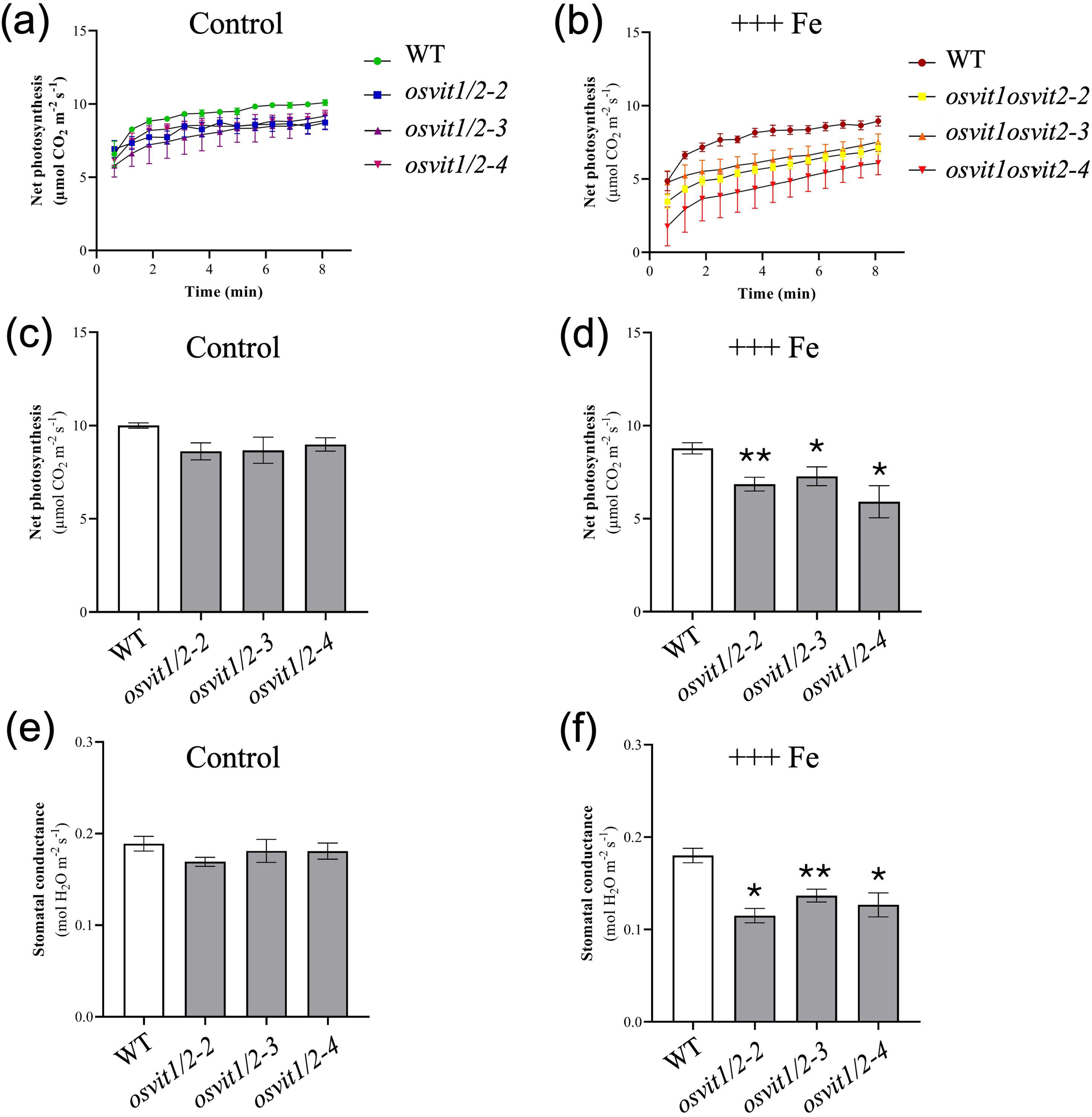
*osvit1osvit2* double knockout mutants net photosynthesis is affected to a larger extent by iron excess. Net photosynthesis curves of WT and *osvit1osvit2* mutants under control (100 µM Fe) (A) and iron excess (5 mM Fe excess) (B) conditions. Net photosynthesis of WT and *osvit1osvit2* mutants under control (C) and iron excess (D) conditions. Stomatal conductance of WT and *osvit1osvit2* mutants under control (E) and iron excess (F) conditions. Asterisks denote statistical significance (* *P* ≤ 0.05; ** *P* ≤ 0.01).

### Mutations in *OsVIT1* and *OsVIT2* genes do not alter leaf and root ionomes

We analyzed the leaf and root ionomes of WT and *osvit1osvit2* plants subjected to control and 5 mM Fe excess for seven days. We focused on consistent concentration differences when comparing WT and mutant plants, as these would suggest a role for VITs in elemental concentration. Although Fe concentration increased in leaves of all Fe excess treated plants compared to control conditions (∼10-fold - Figure 4a and 4b), there was no consistent difference in concentration (*i.e.*, increase or decrease in all mutants) when comparing mutants and WT plants (Fig 4a). In roots, the increase in Fe concentration was even more pronounced (∼15-25-fold), as expected. However, once again, there was no consistent change in Fe concentration when comparing mutants and WT plants under the same condition (Fig 4c and 4d). These data indicate that *osvit1osvit2* double knockout does not affect Fe concentrations at the whole tissue level when compared to WT, either under control or Fe excess conditions. Therefore, the decreased Fe excess tolerance in *osvit1osvit2* plants is likely due to changes in Fe compartmentalization, rather than changes in bulk concentration. We also did not observe differences in concentrations of other elements when comparing mutants and WT plants (Figure S4).

**Figure 4.**
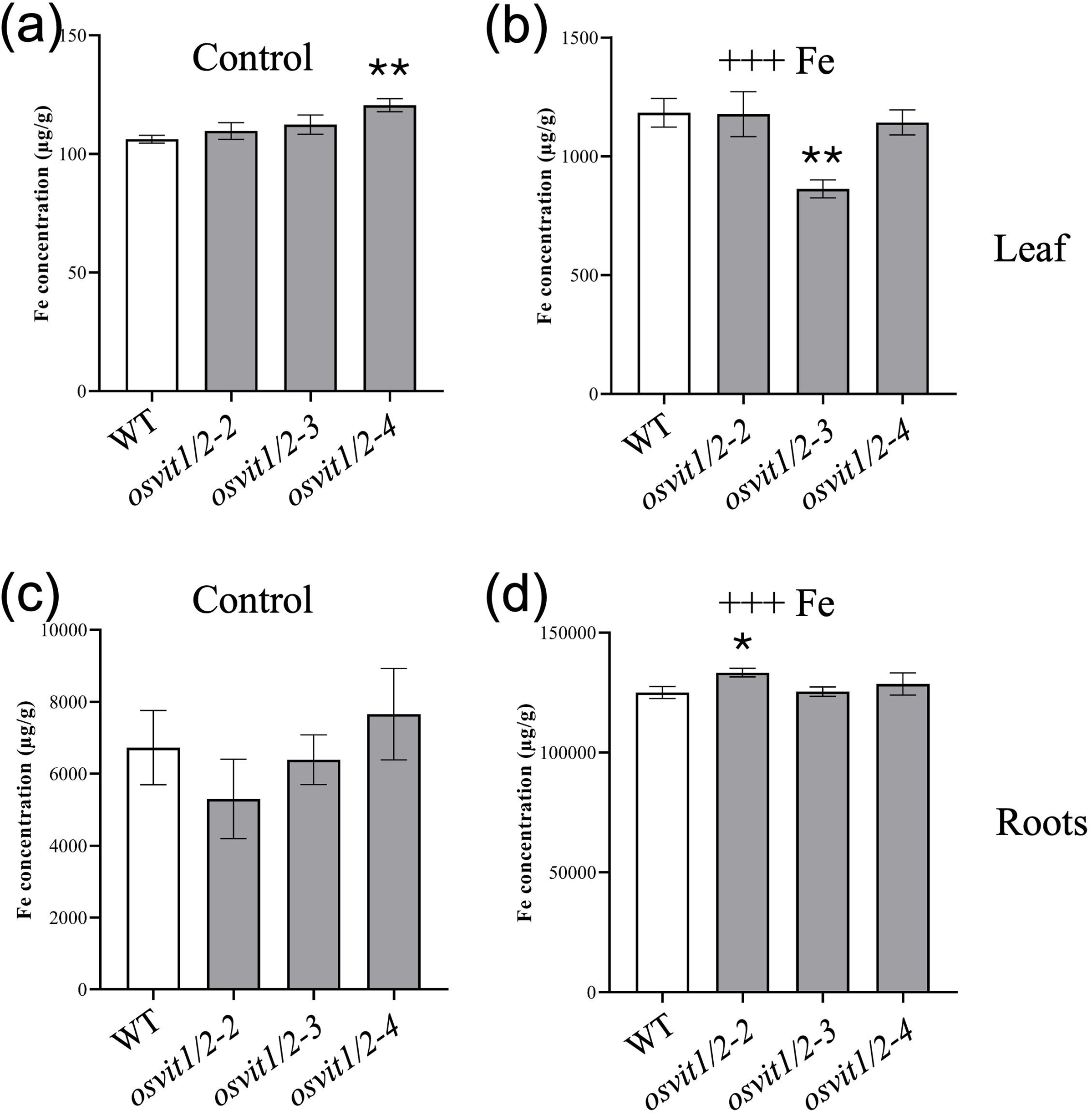
*osvit1osvit2* double knockout mutants have no differences in leaf and root iron concentration in control and iron excess conditions compared to WT. Fe concentration in leaves of WT and *osvit1osvit2* mutants under (A) control and (B) Fe excess (5 mM). Fe concentration in roots of WT and *osvit1osvit2* mutants under (C) control and (D) Fe excess (5 mM). Asterisks denote statistical significance (* *P* ≤ 0.05; ** *P* ≤ 0.01).

### *osvit1osvit2* mutant plants have altered seed ionome

Given the previous reports that single mutants *osvit1* or *osvit2* increased Fe concentration in seeds (Zhang *et al*., 2012; Che *et al*., 2021), we analyzed seed ionomic profiles by ICP-MS. We conducted two independent experiments: the first included WT, *osvit1osvit2*-*1*, and *osvit1osvit2*-*2*; the second included WT, *osvit1osvit2-2*, *osvit1osvit2*-*3*, *osvit1osvit2*-*4*, and *osvit1osvit2*-*5*. Clearly, Fe concentration in non-polished, brown rice was higher in all five mutant lines compared to WT, with a percentage increase ranging from 30% to 100% (Fig 5a and 5b). Similarly, Zn concentrations were also higher in mutant lines (Figure S5a). No consistent changes were observed in the concentration of other elements (Figure S5b-h). We observed that lines with decreased seed filling tended to show increased concentrations of several elements, likely due to a concentration effect (Figure S6). Nonetheless, Fe concentration remained consistently higher in mutant lines compared to WT, and even lines without substantial changes in seed filling exhibited increased Fe levels. Together, these results suggest that the simultaneous mutation of *OsVIT1* and *OsVIT2* can increase Fe and Zn concentrations in rice seeds.

**Figure 5.**
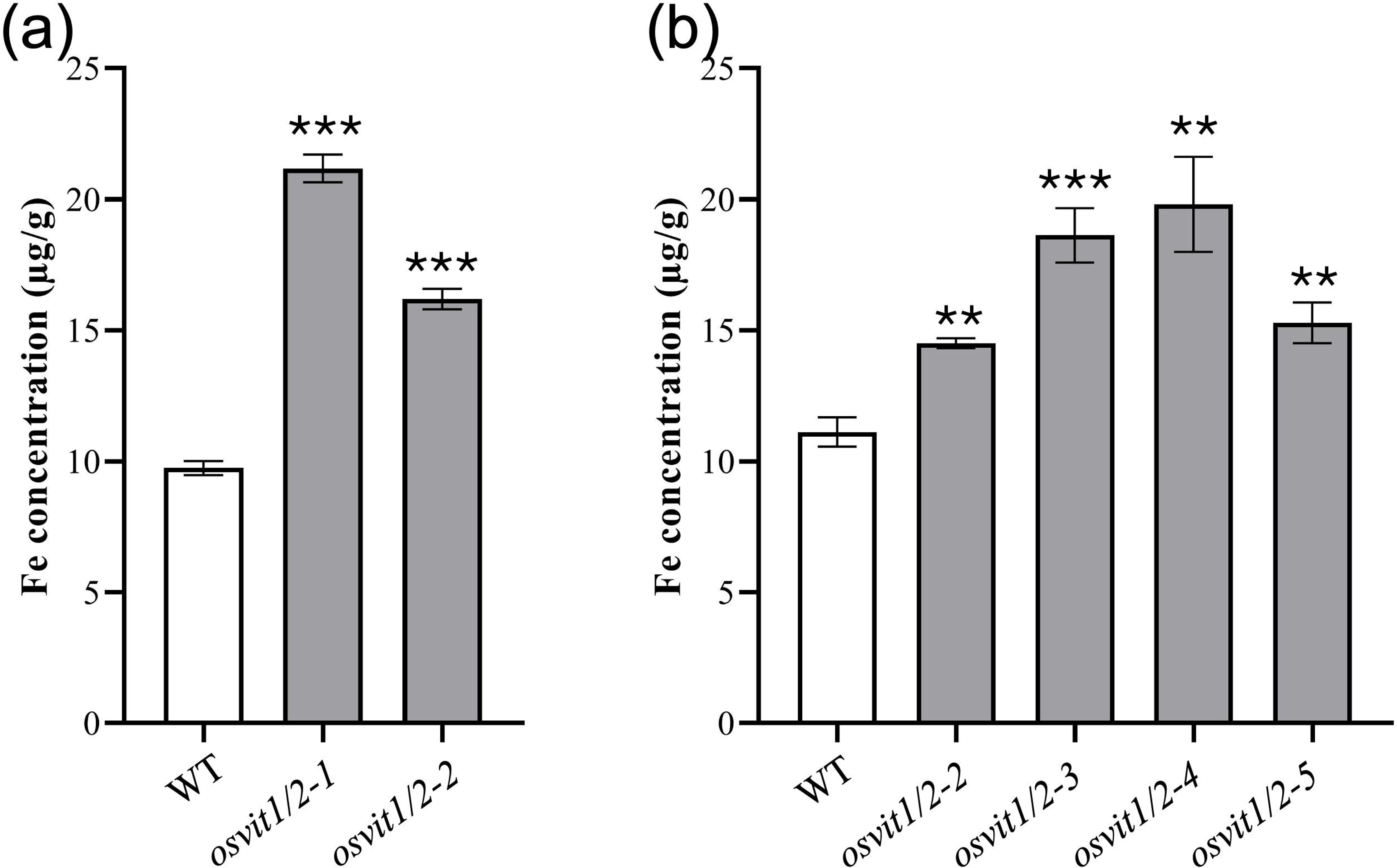
*osvit1osvit2* double knockout mutants show increased iron concentration in seeds compared to WT. (A) Fe concentration in seeds of WT, *osvit1osvit2-1* and *osvit1osvit2-2* plants. (B) Fe concentration in seeds of WT, *osvit1osvit2-2*, *osvit1osvit2-3*, *osvit1osvit2-4* and *osvit1osvit2-5* plants. Asterisks denote statistical significance (* *P* ≤ 0.05; ** *P* ≤ 0.01; *** *P* ≤ 0.001).

### Increased Fe concentration in *osvit1osvit2* seeds is independent of external Fe supply

Next, we investigated whether the increased Fe concentration in seeds of *osvit1osvit2* plants was dependent on external Fe supply. To test this, we cultivated plants from WT, *osvit1osvit2*-*1*, and *osvit1osvit2*-*2* genotypes in a hydroponic system. At anthesis (R4 stage - (Counce *et al*., 2000)), plants were transferred to control conditions, Fe-deficient media (no Fe added), or high-Fe media (1 mM Fe as FeSO_4_) until seed set (R9 stage - (Counce *et al*., 2000)). Flag leaves and seeds were collected and analyzed by ICP-MS. The Fe concentration in flag leaves varied according to the Fe treatment but showed no consistent differences when comparing *osvit1osvit2* plants to WT in all treatments (Fig 6a-c). Interestingly, both *osvit1osvit2* mutant lines showed increased seed Fe concentration compared to WT, regardless of the Fe supply (Fig 6d-f). Taken together, these results indicate that the higher Fe concentration observed in seeds of *osvit1osvit2* mutant plants is independent of external Fe supply.

**Figure 6.**
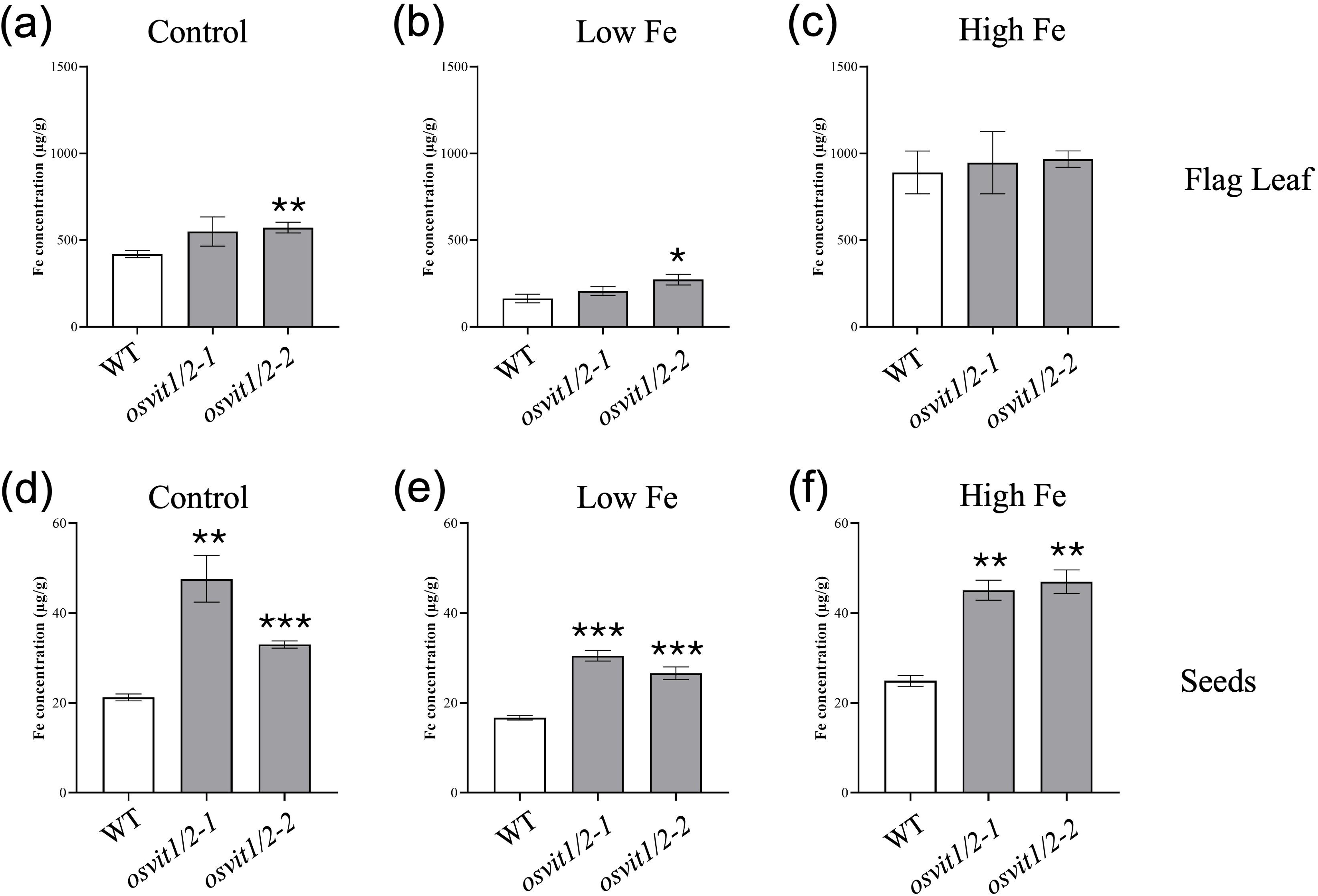
*osvit1osvit2* double knockout mutants increased iron concentration in seeds is independent of external iron supply. (A) Fe concentration in flag leaves of WT, *osvit1osvit2-1* and *osvit1osvit2-2* plants cultivated under control conditions. (B) Fe concentration in flag leaves of WT, *osvit1osvit2-1* and *osvit1osvit2-2* plants cultivated under Fe deficiency after anthesis. (C) Fe concentration in flag leaves of WT, *osvit1osvit2-1* and *osvit1osvit2-2* plants cultivated under high Fe (1 mM) conditions after anthesis. (D) Fe concentration in seeds of WT, *osvit1osvit2-1* and *osvit1osvit2-2* plants cultivated under control conditions. (E) Fe concentration in seeds of WT, *osvit1osvit2-1* and *osvit1osvit2-2* plants cultivated under Fe deficiency after anthesis. (F) Fe concentration in seeds of WT, *osvit1osvit2-1* and *osvit1osvit2-2* plants cultivated under high Fe (1 mM) conditions after anthesis. Asterisks denote statistical significance (* *P* ≤ 0.05; ** *P* ≤ 0.01; *** *P* ≤ 0.001).

### Root-derived Fe contributes to increased Fe in *osvit1osvit2* seeds

Both *OsVIT1* and *OsVIT2* genes are expressed in flag leaves and have been suggested to play a role in Fe compartmentalization in leaf vacuoles, reducing availability for phloem translocation during seed filling (Zhang *et al*., 2012). To test whether the increased Fe concentration in seeds was primarily derived from flag leaves or roots, we supplied stable isotope ^57^Fe to roots or to flag leaves through cuttings. Two concentrations were used for flag leaves: 10 and 100 µM; and 10 µM concentration for root feeding. The concentration of natural ^56^Fe isotope was higher in seeds of *osvit1osvit2* compared to WT, regardless of whether extra ^57^Fe was supplied through flag leaves or roots (Fig 7a-c). However, ^57^Fe concentration was significantly higher in seeds of the mutant only when ^57^Fe was supplied to roots (Fig 7d-f). These results suggest that Fe uptake and translocation from roots contribute to the elevated Fe concentration in seeds of *osvit1osvit2* plants. On the other hand, our results did not show a clear contribution of flag leaf-derived Fe to the increased Fe concentration in seeds of *osvit1osvit2* plants.

**Figure 7.**
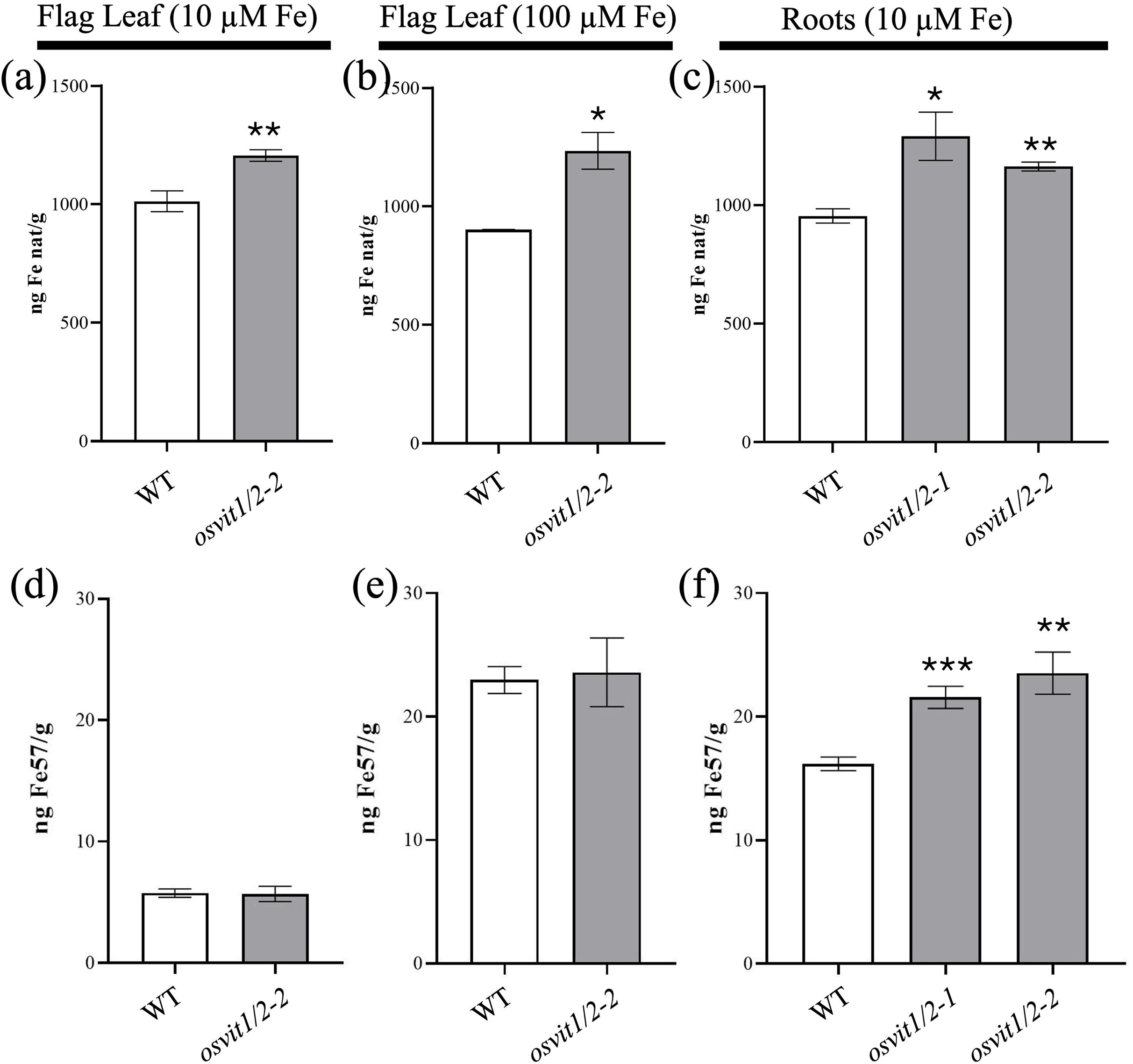
Root uptake is important for iron increase in *osvit1osvit2* double knockout mutants. ^56^Fe (natural Fe) quantification in seeds of plants which had their flag leaves fed with 10 µM Fe (A), 100 µM Fe (B); or their roots fed with 10 µM Fe (C). ^57^Fe isotope quantification in seeds of plants which had their flag leaves fed with 10 µM Fe (D), 100 µM Fe (E); or their roots fed with 10 µM Fe (F). Asterisks denote statistical significance (* *P* ≤ 0.05; ** *P* ≤ 0.01; *** *P* ≤ 0.001).

### Seeds of *osvit1osvit2* plants have altered Fe distribution

It was previously reported that the spatial distribution of Fe in seeds of *osvit1* and *osvit2* mutants is altered (Zhang *et al*., 2012). Specifically, both *osvit1* and *osvit2* T-DNA mutants show increased Fe accumulation in the embryo (Zhang *et al*., 2012), whereas *osvit2* gene-edited plants have higher Fe in the embryo and lower Fe in the aleurone layer (Che *et al*., 2021). Besides these observations, differences in elemental distribution within seeds of those genotypes were not explored in detail

To determine whether the double mutant lines exhibited changes in Fe distribution within seeds, we used X-ray fluorescence spectroscopy (XRF) to scan across seed sections. We observed that *osvit1osvit2* double mutants displayed increased Fe levels in the endosperm compared to WT (Fig 8a and 8b). Next, we employed synchrotron-based X-ray fluorescence spectroscopy (SXRF) to map the spatial distribution of longitudinal sections of embryos. The embryos of *osvit1osvit2* double mutants exhibited elevated Fe concentrations and altered Fe distribution compared to WT, with greater Fe accumulation throughout the embryo (Fig 9a). Furthermore, we did not detect a clear difference in Zn distribution (Fig 9b), while Mn distribution seemed altered, with increased accumulation in the shoot apical meristem of *osvit1osvit2* double mutant seeds (Fig 9c).

**Figure 8.**
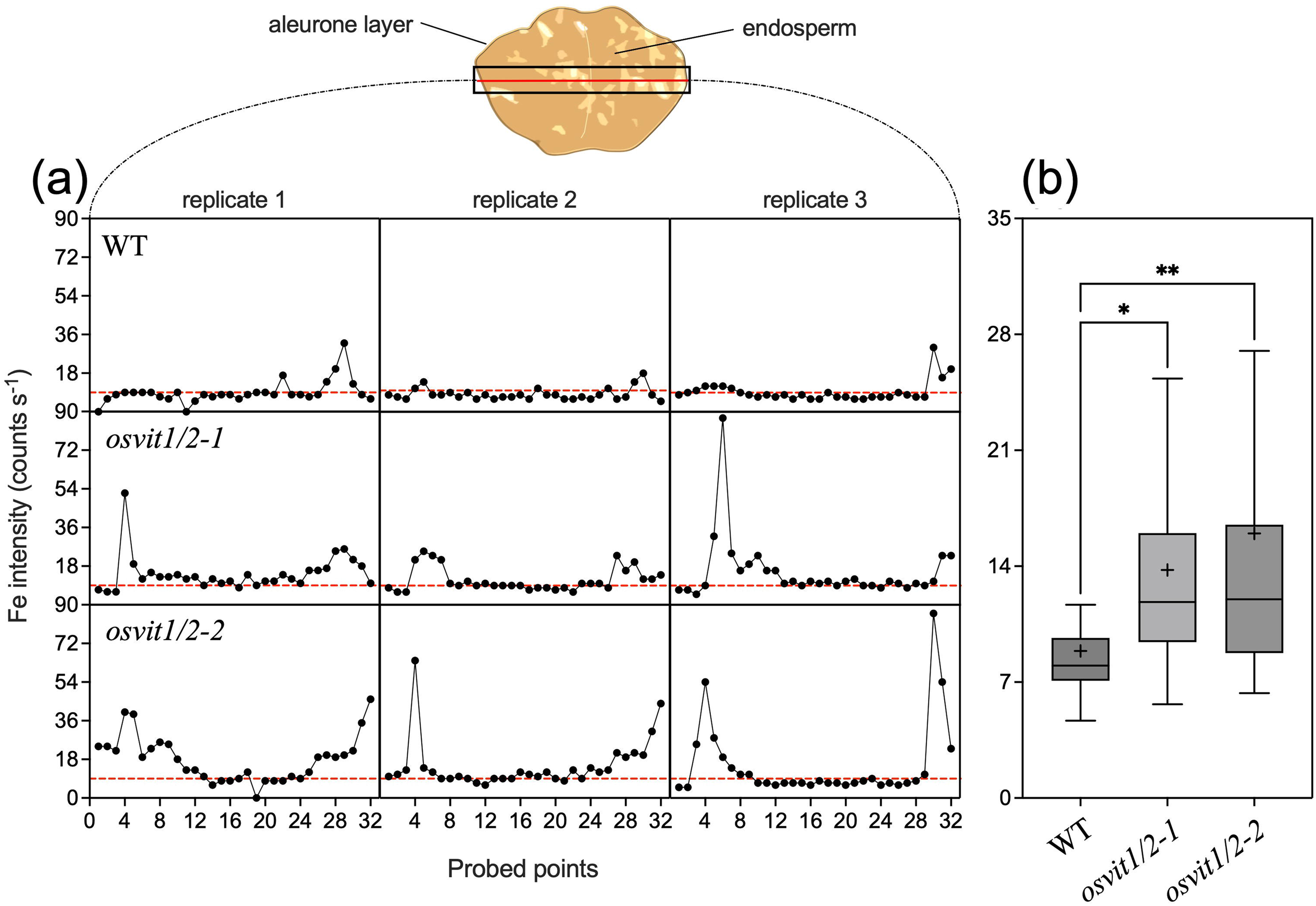
Iron accumulates in the endosperm of *osvit1osvit2* double knockout mutants. (A) Linescans of WT, *osvit1osvit2-1* and *osvit1osvit2-2* mutant seeds cross sections. Three replicates per genotype are shown. Each graph showed all points measured across the section. The red line denotes the average of measurements of WT. (B) Comparison of all points measured in each genotype. Boxplots show the first and third percentile; lines show the median; crosses show the average; and bars show the minimum and maximum values. Asterisks denote statistical significance (* *P* ≤ 0.05; ** *P* ≤ 0.01).

**Figure 9.**
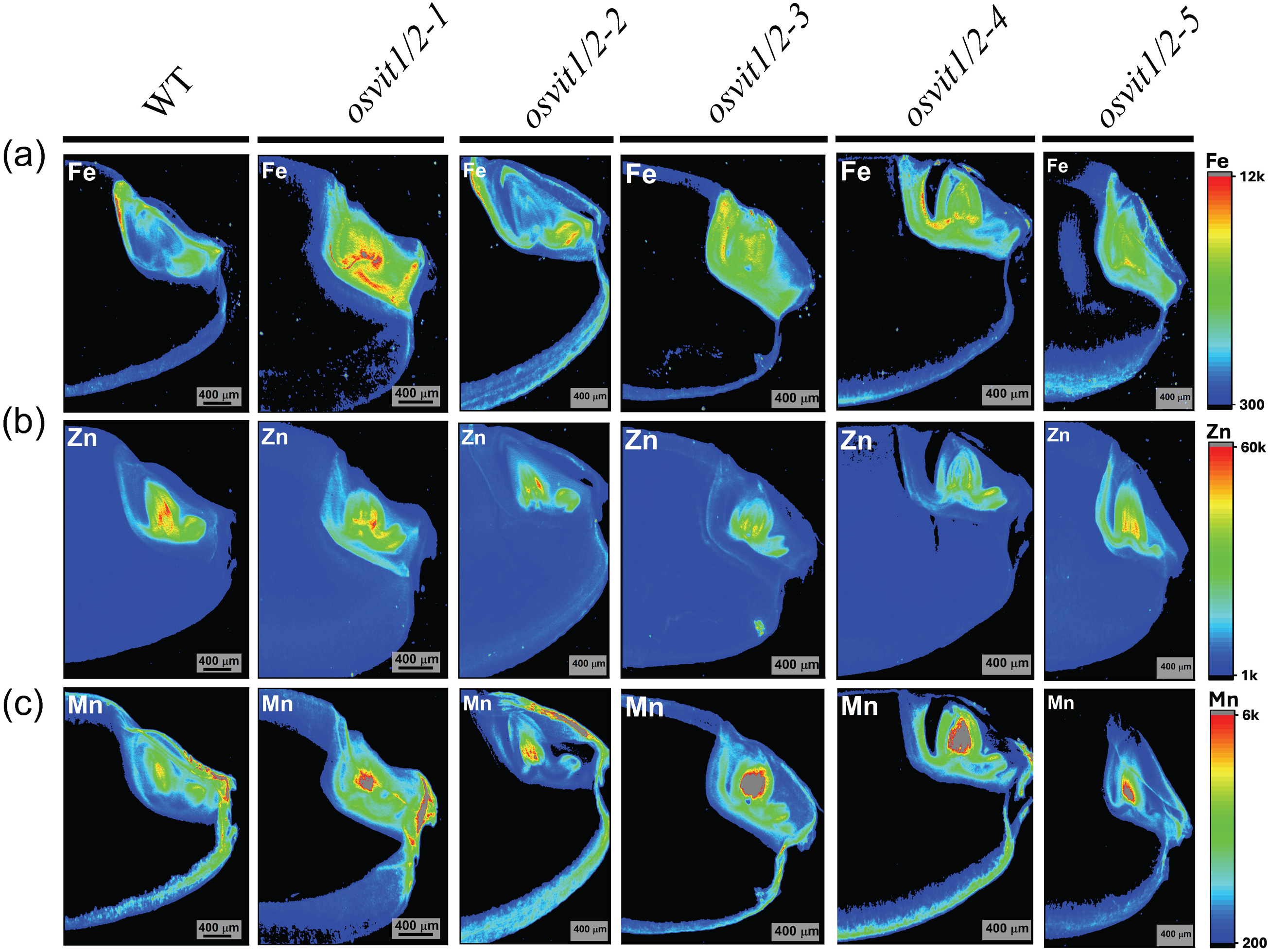
Synchrotron X-ray fluorescence shows that *osvit1osvit2* double knockout mutants have altered iron distribution in the embryo. (A) Fe distribution in seeds of WT and five *osvit1osvit2-2* mutants. (B) Zn distribution in seeds of WT and five *osvit1osvit2-2* mutants. (C) Mn distribution in seeds of WT and five *osvit1osvit2-2* mutants. Data were acquired at BioXAS beamline of the Canadian Light Source (CLS, Saskatoon, Canada).

We further examined the spatial distribution of elements through SXRF at a beamline that allowed higher resolution to map WT and *osvit1osvit2-2* embryos. Using signal intensity to compare elemental distribution and abundance, we observed that *osvit1osvit2-2* embryos exhibited a distinct Fe distribution pattern as well as increased Fe abundance compared to WT (Fig 10a). While in WT Fe mainly accumulated in the radicle and scutellum, *osvit1osvit2-2* embryos accumulated Fe to a larger extent in the scutellum and presented Fe in the plumule (Fig 10a and 10b; please see Figure S7 for reference on regions mapped at higher resolution). The distributions of Zn and Mn, on the other hand, are more similar in both genotypes, although *osvit1osvit2-2* embryos seem to accumulate more Mn in the scutellum and mainly in the plumule (Fig 10a and 10b; also see Fig 9c). When the localization of Fe, Zn and Mn were compared in the same maps, the increased accumulation and altered distribution of Fe was also obvious (Fig 10c and 10b). Interestingly, Fe accumulation in the scutellum of and plumule of *osvit1osvit2-2* overlaps that of Zn (Fig 10d).

**Figure 10.**
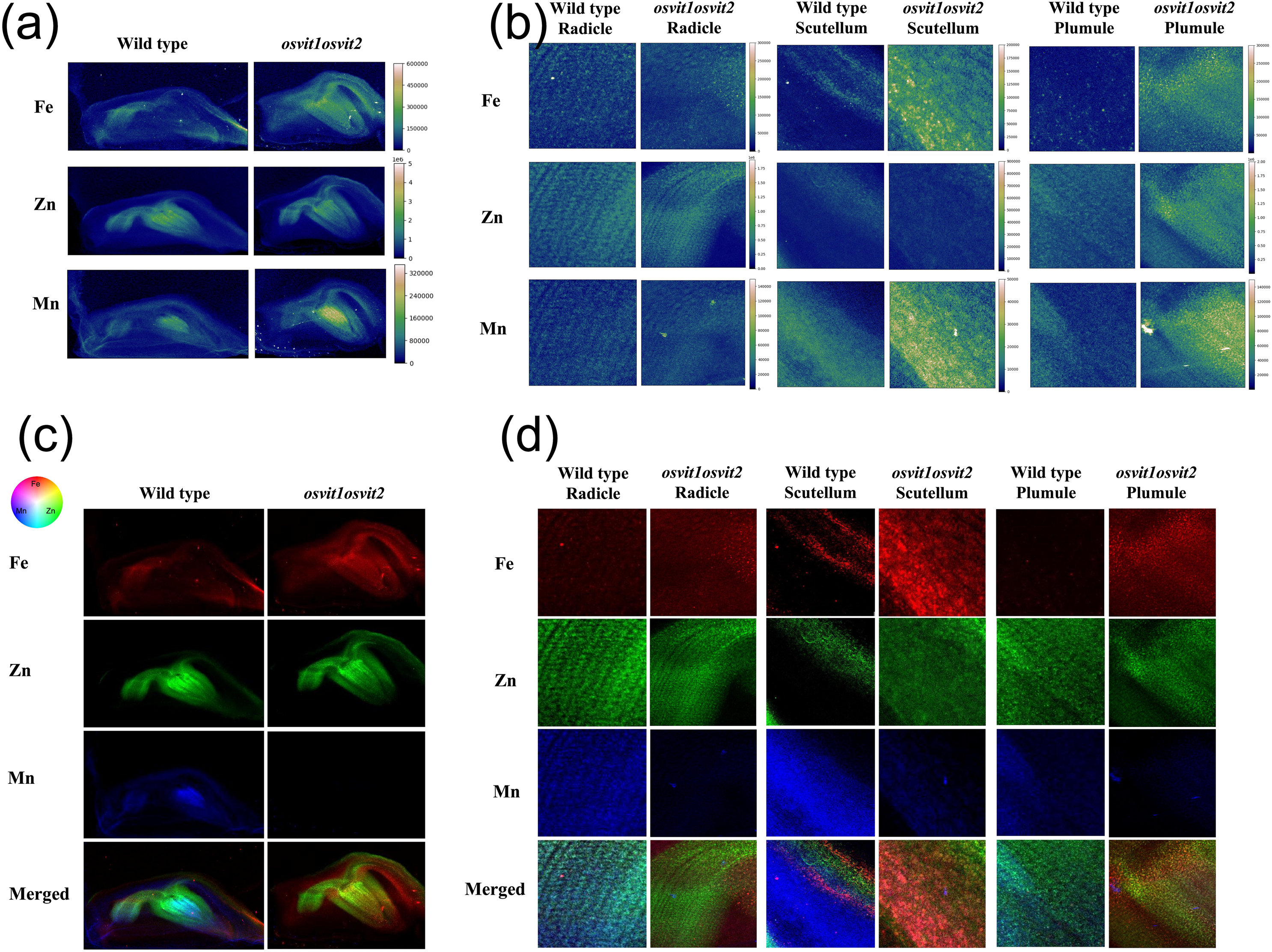
Embryos from *osvit1osvit2* double knockout mutants have altered iron distribution and increased iron accumulation in scutellum and plumule. (A) Panoramic maps showing signal intensity of Fe, Zn and Mn distribution in embryos of WT and *osvit1osvit2-2*. Scale is the same for both genotypes. (B) High resolution maps showing signal intensity of Fe, Zn and Mn distribution in radicles, scutellum and plumule of WT and *osvit1osvit2-2*. Scale is the same for both genotypes. (C) Panoramic maps comparing Fe (red), Zn (green) and Mn (blue) distribution in RGB in embryos of WT and *osvit1osvit2-2*. (D) Low resolution maps comparing Fe (red), Zn (green) and Mn (blue) distribution in RGB in radicles, scutellum and plumule of WT and *osvit1osvit2-2*. Data were acquired at CARNAUBA Beamline at SIRIUS, Laboratório Nacional de Luz Síncrotron/Centro Nacional de Energia e Materiais (LNLS/CNPEM, Campinas, SP, Brazil).

To further investigate how Fe distribution is altered in *osvit1osvit2* mutants, we performed Perls/DAB staining on thin seed sections from *osvit1osvit2* mutants and WT plants. Our analysis revealed that scutellum cells at the abaxial surface in all five mutant lines accumulated more Fe than WT (Fig 11a-f). The Fe staining exhibited a punctuated pattern surrounding the nuclei of the subepidermal cells (Fig 11a-f). However, we did not observe any alteration in Fe distribution within the aleurone layer (Figure S8). Altogether, these findings suggest that increased Fe accumulation in the scutellum and plumule contributes to the Fe enrichment in *osvit1osvit2* mutant seeds.

**Figure 11.**
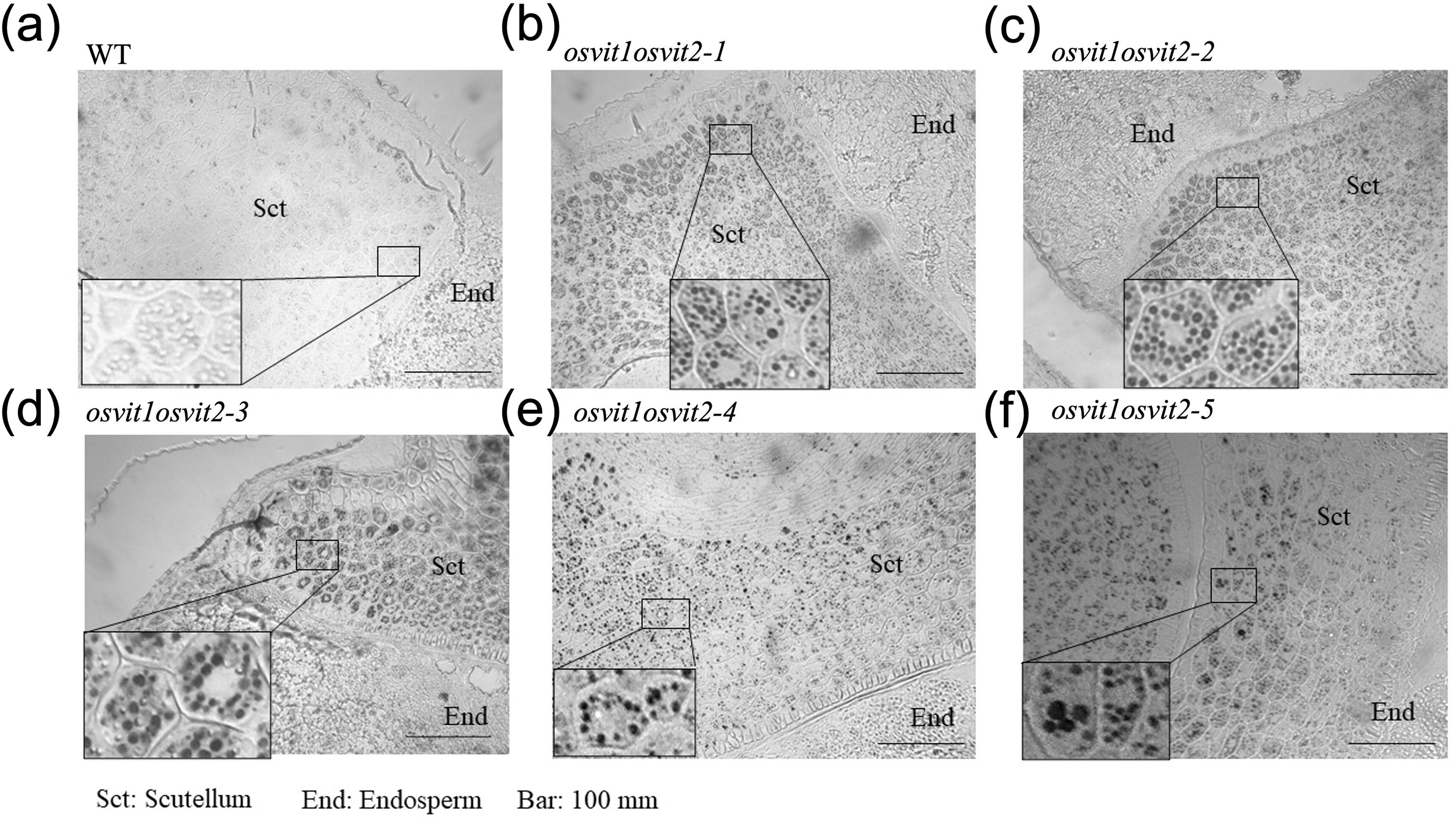
Iron staining with Perls/DAB shows higher accumulation in the scutellum in a punctuated pattern. (A) WT; (B) *osvit1osvit2-1*; (C) *osvit1osvit2-2*; (D) *osvit1osvit2-3*; (E) *osvit1osvit2-4*; (F) *osvit1osvit2-4*. Bars = 100 µm.

## Discussion

### Mutations in *VIT* genes decrease Fe excess tolerance in rice

Fe toxicity is a significant constraint for rice cultivation under specific environmental conditions, such as waterlogged, poorly drained, or acidic soils. Since most plants do not face Fe excess in their natural environments, it is likely that mechanisms for Fe excess tolerance are not as ancient and phylogenetically conserved as those for Fe deficiency (Chao & Chao, 2022; Rodrigues *et al*., 2023). Nevertheless, general strategies for Fe excess tolerance in rice have been proposed based on whether different genotypes exclude Fe at the root level (Defense I), compartmentalize Fe in roots to prevent translocation to shoots (Defense II), compartmentalize Fe within shoots (Defense III), or detoxify ROS generated by Fe overload (Defense IV) (Aung & Masuda, 2020; Wairich *et al*., 2024). Despite these proposed strategies, only a few genes directly involved in Fe excess tolerance have been characterized to date. In this study, we demonstrated that rice plants lacking functional VIT transporters exhibit reduced tolerance to Fe excess.

Several transcriptomic and RT-qPCR analyses have shown that *OsVIT2* is up-regulated in both roots and shoots in response to Fe excess (Zhang *et al*., 2012; Finatto *et al*., 2015; Stein *et al*., 2019; Wairich *et al*., 2021; Regon *et al*., 2022). Both OsVIT1 and OsVIT2 can complement *ccc1* yeast mutants (Zhang *et al*., 2012; Che *et al*., 2021), indicating they are functional vacuolar Fe transporters. These findings suggest that VITs, particularly OsVIT2, play a role in Fe detoxification under conditions of excess Fe availability. A previous study tested an *osvit2* mutant line carrying a T-DNA insertion upstream of the coding sequence, resulting in *OsVIT2* knockdown, and observed slightly shorter shoots under Fe excess (Bashir *et al*., 2013). However, *osvit1* and *osvit2* single mutants analyzed by Zhang *et al*. (2012) showed no difference from WT under high Fe conditions. Other studies that identified mutants for rice *VIT* genes have not reported on Fe excess tolerance. Thus, there is no previous evidence that single *osvit1* or *osvit2* mutants exhibit reduced tolerance to Fe excess (Zhang *et al*., 2012; Bashir *et al*., 2013; Che *et al*., 2021; Kandwal *et al*., 2022).

In contrast, our *osvit1osvit2* double mutant lines were clearly less tolerant to excessive Fe compared to WT (Fig 2). We observed deleterious effects on photosynthesis (Fig 3), demonstrating that loss of function of *VIT* genes decreases the capacity to tolerate high Fe concentrations. However, *osvit1* or *osvit2* single mutants have not been reported as less tolerant (Zhang *et al*., 2012; Bashir *et al*., 2013). This discrepancy could be attributed to (1) distinct experimental conditions among studies or (2) partial redundancy of OsVIT1 and OsVIT2 in Fe detoxification. Our experiments were conducted at 5 mM Fe concentration, which is considerably higher than those used in previous studies (0.5 mM and 1 mM; Bashir *et al*., 2013 and Zhang *et al*., 2012, respectively). Under our growth conditions, Fe concentrations up to 1 mM are not toxic to rice plants, causing little to no leaf bronzing. Therefore, it is possible that single mutants also exhibit reduced tolerance compared to WT but were not tested under conditions severe enough to reveal this phenotype. *OsVIT1* and *OsVIT2* are not entirely redundant, as their expression is differentially regulated under Fe excess, and single mutations lead to different seed phenotypes, indicating they do not fully compensate for each other (Zhang *et al*., 2012). Thus, it is also plausible that only double mutants exhibit sensitivity to Fe excess, as both OsVIT1 and OsVIT2 function as vacuolar Fe importers and may act partially redundantly in Fe detoxification.

Interestingly, the lower tolerance observed in *osvit1osvit2* double mutant lines does not stem from alterations in Fe concentration in leaves or roots, as mutants and WT plants exhibited similar Fe levels (Fig 4). This suggests that the primary cause of decreased tolerance is the inability to detoxify Fe within vacuoles, leading to toxicity when intracellular Fe distribution is disrupted. Our data also indicate that leaf bronzing (Fig 2) likely results from impaired vacuolar Fe detoxification in leaf cells. Both *VIT* genes are expressed in roots, with *OsVIT2* being induced under Fe excess conditions (Zhang *et al*., 2012; Che *et al*., 2021). Consequently, the loss of root vacuolar transporters would be expected to increase Fe concentration in aerial tissues of *osvit1osvit2* plants under control or high Fe conditions, similar to how other vacuolar root transports function (Huang *et al*., 2016; Pita-Barbosa *et al*., 2019). However, compensatory mechanisms, such as the activity of alternative vacuolar Fe transporters, may compensate for Fe sequestration in roots, preventing significant Fe accumulation in shoots. As a result, phenotypic alterations are primarily observed in leaves of *osvit1osvit2* plants exposed to Fe excess, likely due to their reduced capacity for Fe detoxification.

### OsVIT1 and OsVIT2 as targets for biofortification

Previous studies have identified *OsVIT1* and *OsVIT2* as potential target genes for biofortification, as independent loss of function of either transporter leads to increased Fe concentration (Zhang *et al*., 2012; Bashir *et al*., 2013; Che *et al*., 2021; Kandwal *et al*., 2022). In this study, we demonstrated that *osvit1osvit2* double mutants recapitulate this phenotype (Fig 5). We also observed that Fe accumulated in embryo, with markedly changes in distribution in the scutellum and the plumule (Fig 9 and Fig 10). Moreover, we identified that Fe accumulation in the scutellum occurs in a punctuated pattern (Fig 11). Overall, our findings indicate that the *osvit1osvit2* mutants display comparable phenotypic traits to those of single *vit* mutants, while demonstrating a role of VIT transporters in Fe homeostasis and their potential as targets for biofortification.

The Fe concentration increases observed in *osvit1osvit2* seeds were similar in magnitude to those reported for *osvit1* or *osvit2* single mutants (Zhang *et al*., 2012; Bashir *et al*., 2013; Che *et al*., 2021; Kandwal *et al*., 2022). This suggests that while the mutations significantly enhance Fe levels compared to WT, the effects of disrupting both *VIT* genes are not additive regarding seed Fe concentration. Consequently, there may be no clear advantage in generating double mutants for biofortification, especially considering the tolerance to high Fe is decreased in *osvit1osvit2* plants (Fig 2). Nonetheless, our findings further reinforce that *VIT* genes are strong candidates for Fe biofortification in rice, as non-transgenic lines can be developed with substantial Fe accumulation in seeds, outperforming several transgenic-based approaches (Wairich *et al*., 2022). Future studies should also consider the impact of combining *OsVIT* mutations with other candidate genes, either by gene editing or transgenic strategies, as single and double *osvit* mutants may yield different phenotypes depending on the biofortification approach.

Previous studies suggested that OsVIT1 and OsVIT2 roles are to sequester Fe in flag leaf vacuoles and that their loss of function would allow more remobilization to developing seeds, which would explain the increased Fe accumulation in grains (Zhang *et al*., 2012). However, our ^57^Fe feeding experiments to either flag leaves or roots demonstrated that root-derived rather than flag-leaf-derived Fe, is responsible for the increased Fe accumulation observed in *osvit1osvit2* mutants (Fig 7). This finding suggests that OsVIT1 and OsVIT2 may function similarly to other vacuolar transporters to limit root-to-shoot translocation and shoot accumulation of their substrates (Arrivault *et al*., 2006; Chao *et al*., 2012; Eroglu *et al*., 2016; Huang *et al*., 2016; Ricachenevsky *et al*., 2018). Since *OsVIT2* is expressed in other tissues such as nodes, which are involved in the vascular nutrient transfer (Che *et al*., 2021), the loss of *OsVIT1* and *OsVIT2* may also decrease Fe restriction at these sites. Given that nodes serve as central hubs for metal accumulation and redistribution (Yamaji & Ma, 2014), their altered function in *osvit1osvit2* plants may contribute to enhanced Fe allocation to seeds. Thus, our results indicate that root Fe uptake plays a role in Fe accumulation in the seeds of *osvit1osvit2* mutant lines, whereas direct remobilization from flag leaves appears to be less relevant under our experimental conditions. Given that Fe remobilization varies depending on Fe availability (Sperotto, 2013), this could explain the observed differences. Furthermore, we demonstrated that the increased Fe accumulation in double mutants occurs independently of external Fe supply (Fig 6), suggesting that the phenotype is stable across varying Fe conditions. Consequently, we expect that the enhanced Fe accumulation in seeds of *osvit1osvit2* mutants will be maintained under diverse growing environments with different Fe availability.

Rice biofortification efforts primarily target the endosperm, the starch-rich central part of the seed that remains in white rice, the most widely consumed form (Wairich *et al*., 2022). Previous studies have shown that *osvit2* mutants display lower Fe in the aleurone layer, the external tissue around the endosperm, suggesting that OsVIT2 sequesters Fe in vacuoles of these cells, while loss of function of *osvit2* mutants bypasses Fe into the endosperm tissue (Che *et al*., 2021). Similarly, our XRF data suggest that *osvit1osvit2* mutants increase Fe accumulation within the endosperm (Fig S8); however, we did not observe changes in Fe accumulation in the aleurone layer (Figure 8). Therefore, whether this effect is restricted to single *osvit2* mutants or also applies to *osvit1osvit2* remains to be determined.

Finally, *osvit1osvit2* mutants showed an altered Fe distribution pattern in embryos (Fig 9), with pronounced Fe accumulation in the scutellum and plumule (Fig 10). Similar changes in embryo Fe distribution were previously reported for single *osvit1* and *osvit2* mutants (Zhang *et al*., 2012; Che *et al*., 2021). XRF data indicate that while Fe is predominantly localized in the scutellum in WT seeds (Lu *et al*., 2013), *osvit1osvit2* mutants accumulate Fe in multiple embryo regions beyond the scutellum (Fig 9 and 10). These shifts in Fe spatial distribution suggest that *VIT* genes play a role in precise Fe localization within embryos, resembling that of *AtVIT1* in *A. thaliana* (Kim *et al*., 2006). Therefore, VIT transporters might play dual roles in Fe translocation to seeds as well as Fe distribution within seed tissues. Moreover, our data reveal that Fe accumulates in distinct punctate structures within scutellum adaxial surface cells. The scutellum appears to be the primary Fe storage site in embryos, with mutants exhibiting higher Fe deposition in these cells compared to WT (Fig 11). The punctate Fe accumulation pattern resembles protein bodies found in scutellum and aleurone cells, which often also contain phytate (Prom u thai *et al*., 2008). Further studies are required to elucidate the physiological consequences of Fe mislocalization in this tissue, particularly regarding seed metabolism and Fe bioavailability.

### Conclusion

We showed that *osvit1osvit2* double mutants have increased Fe concentration in seeds, particularly in the embryo, with elevated Fe levels in the coleoptile and scutellum. Our findings indicate that root uptake and root-to-shoot translocation contribute to this Fe accumulation, and that the effect of mutating both *VIT* genes on Fe concentration is more pronounced than the effect of external Fe supply. Furthermore, we established that the lack of both *OsVIT1* and *OsVIT2* genes reduces Fe excess tolerance in rice plants, suggesting that targeting these genes for biofortification could have potential trade-offs in terms of plant fitness in Fe-rich, waterlogged field conditions.

## Supporting information

Figure S1: Sequence alignment of the predicted translated polypeptides of the OsVIT1 and OsVIT2 in all five osvit1osvit2 mutants.

Figure S2: All five osvit1osvit2 double knockout lines are more sensitive to iron excess.

Figure S3: Growth of osvit1osvit2 double knockout lines is not severely affected by iron excess.

Figure S4: Elemental profiling of WT and osvit1osvit2 leaves and roots of plants under control and iron excess.

Figure S5: Elemental profiling of WT and osvit1osvit2 seeds.

Figure S6: osvit1osvit2 double knockout mutants changes in agronomic traits.

Figure S7: Regions from WT and osvit1osvit2 embryos selected for high resolution analyses using Synchrotron X-Ray Fluorescence.

Figure S8: Iron staining with Perls/DAB in aleurone layer cells.

Table S1. Primers used in this work.

## Acknowledgements

This work was financed by Conselho Nacional de Desenvolvimento Científico e Tecnológico (CNPq; grants numbers 442304/2023-4, 444596/2024-0), Fundação de Amparo à Pesquisa do Estado do Rio Grande do Sul (FAPERGS; grants number 21/2551-0001960-3, 23/2551-0001862-4 and 22/2551-0000834-8) and Instituto Serrapilheira (grant number 1709-17256). This study was financed in part by the Coordenação de Aperfeiçoamento de Pessoal de Nível Superior - Brasil (CAPES) - Finance Code 001.

## Author contributions

FKR and FSM conceived the study. BDB, AGSR, RVO, FKR, FSM, SC and TM designed experiments. BDB, AGSR, RVO, YLM, ES, GSM, JSA, LRP, VHRF, FMMB, FO, JPQM, RAS, HWPC, RT, GMB, NN, HR, CAP, RFHG, MMP, FSM and FKR conducted research, performed data collection, analyses and interpretation. FSM and FKR wrote the manuscript. All authors helped improve the manuscript and approved the submitted version.

## Data availability

All data is available upon request.

## Supporting Information

**Figure S1: Sequence alignment of the predicted translated polypeptides of the OsVIT1 and OsVIT2 in all five *osvit1osvit2* mutants.** The predicted full WT sequence of *OsVIT1* and *OsVIT2* is shown on top of each alignment. All mutants present predicted truncated versions of the WT proteins. Shaded aminoacids highlight the WT conserved sequence.

**Figure S2: All five *osvit1osvit2* double knockout lines are more sensitive to iron excess.** (A) Leaves of wild type and five *osvit1osvit2* under control conditions (100 µM Fe). (B) Leaves of wild type and five *osvit1osvit2* were exposed to seven days of 5 mM Fe excess.

**Figure S3: Growth of *osvit1osvit2* double knockout lines is not severely affected by iron excess.** Culm height of WT and *osvit1osvit2* mutants in control (A) and iron excess (B) conditions after seven days treatment. Shoot dry weight of WT and *osvit1osvit2* mutants in control (C) and iron excess (D) conditions. Root length of WT and *osvit1osvit2* mutants in control (E) and iron excess (F) conditions. Root dry weight of WT and *osvit1osvit2* mutants in control (G) and iron excess (H) conditions. SPAD values of WT and *osvit1osvit2* mutants in control (I) and iron excess (J) conditions. Fe concentrations were 100 µM Fe in control and 5 mM in Fe excess. Asterisks denote statistical significance (* *P* ≤ 0.05; ** *P* ≤ 0.01; *** *P* ≤ 0.001).

**Figure S4: Elemental profiling of WT and *osvit1osvit2* leaves and roots of plants under control and iron excess.** (A) Calcium; (B) copper; (C) potassium; (D) magnesium; (E) manganese; (F) phosphorus; (G) sulfur; (H) zinc. Asterisks denote statistical significance (* *P* ≤ 0.05; ** *P* ≤ 0.01; *** *P* ≤ 0.001).

**Figure S5: Elemental profiling of WT and *osvit1osvit2* seeds.** (A) zinc; (B) copper; (C) potassium; (D) magnesium; (E) manganese; (F) phosphorus; (G) sulfur; (H) calcium. Asterisks denote statistical significance (* *P* ≤ 0.05; ** *P* ≤ 0.01; *** *P* ≤ 0.001).

**Figure S6: *osvit1osvit2* double knockout mutants changes in agronomic traits.** (A) 1,000 seeds weight in WT and *osvit1osvit2* mutant lines. (B) Plant height in WT and *osvit1osvit2* mutant lines. (C) Filled seeds, empty seeds and total seeds in WT and *osvit1osvit2* mutant lines. (D) Relative seed filling in WT and *osvit1osvit2* mutant lines. Asterisks denote statistical significance (* *P* ≤ 0.05; ** *P* ≤ 0.01; *** *P* ≤ 0.001). Letter are used to denote differences in total number of seeds comparing WT and *osvit1osvit2* mutants.

**Figure S7:** Regions from WT and *osvit1osvit2* embryos selected for high resolution analyses using Synchrotron X-Ray Fluorescence. (A) Wild type; (B) *osvit1osvit2-2*. Squares show selected regions of radicle, scutellum and plumule in each embryo.

**Figure S8: Iron staining with Perls/DAB in aleurone layer cells.** (A) WT; (B) *osvit1osvit2-1*; (C) *osvit1osvit2-2*; (D) *osvit1osvit2-3*; (E) *osvit1osvit2-4*; (F) *osvit1osvit2-5*. Bars = 100 µm.

