## Supplementary figures and images for "Double knockout of rice *OsVIT1* and *OsVIT2* genes reveals a trade-off between iron biofortification and iron excess tolerance"

### Figure S1: Sequence alignment of the predicted translated polypeptides of the OsVIT1 and OsVIT2 in all five osvit1osvit2 mutants.

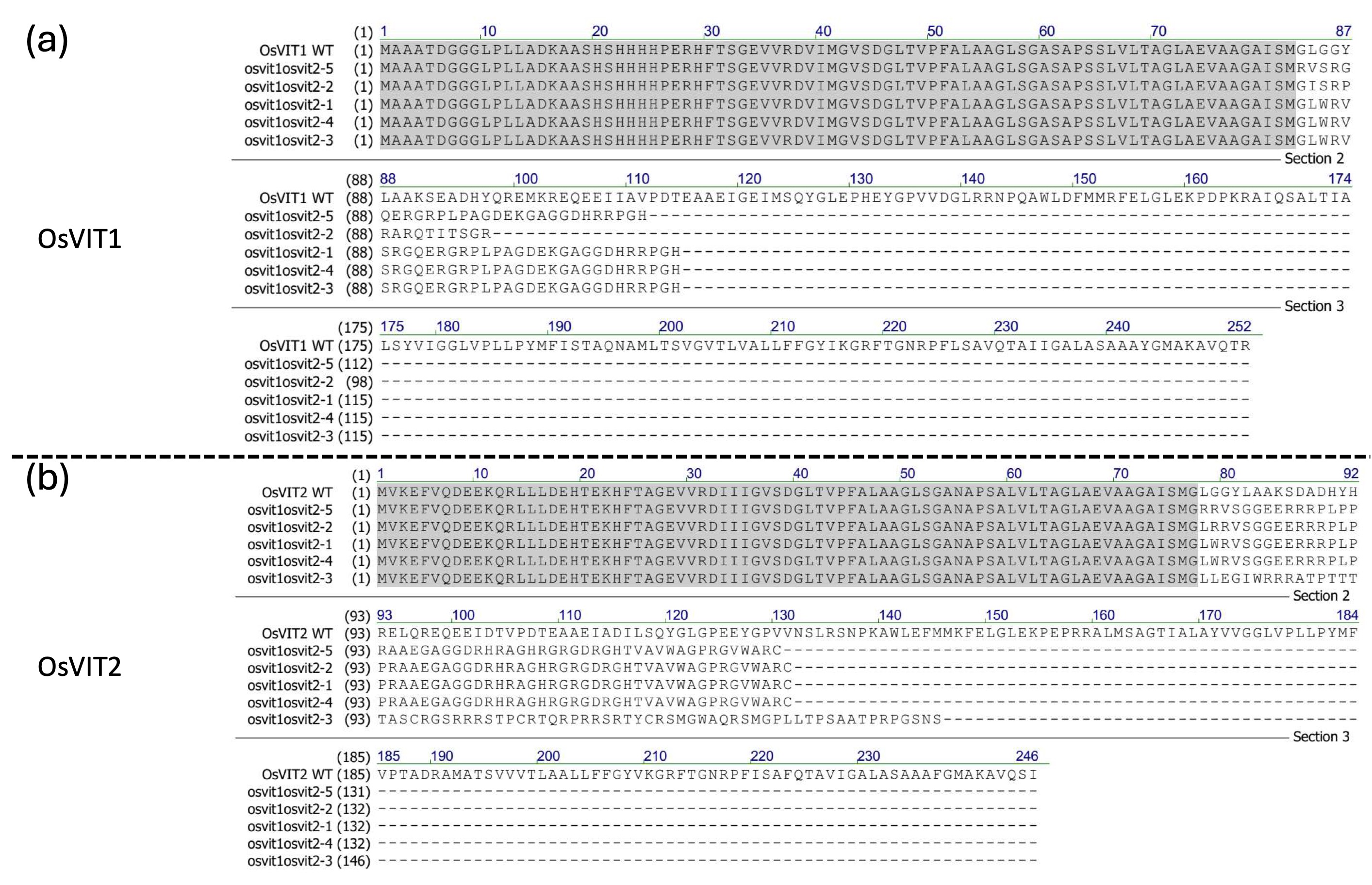

### Figure S2: All five osvit1osvit2 double knockout lines are more sensitive to iron excess.

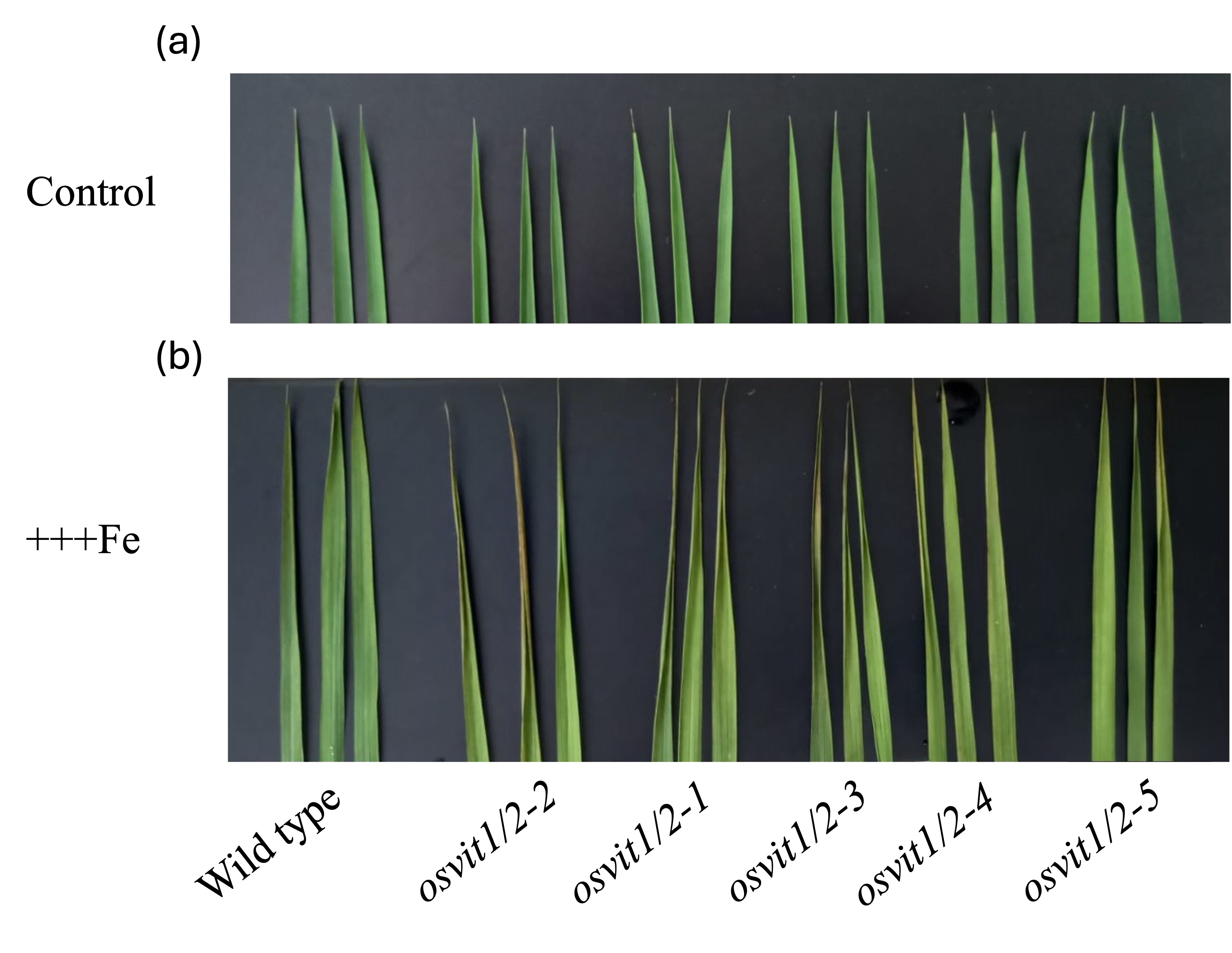

### Figure S3: Growth of osvit1osvit2 double knockout lines is not severely affected by iron excess.

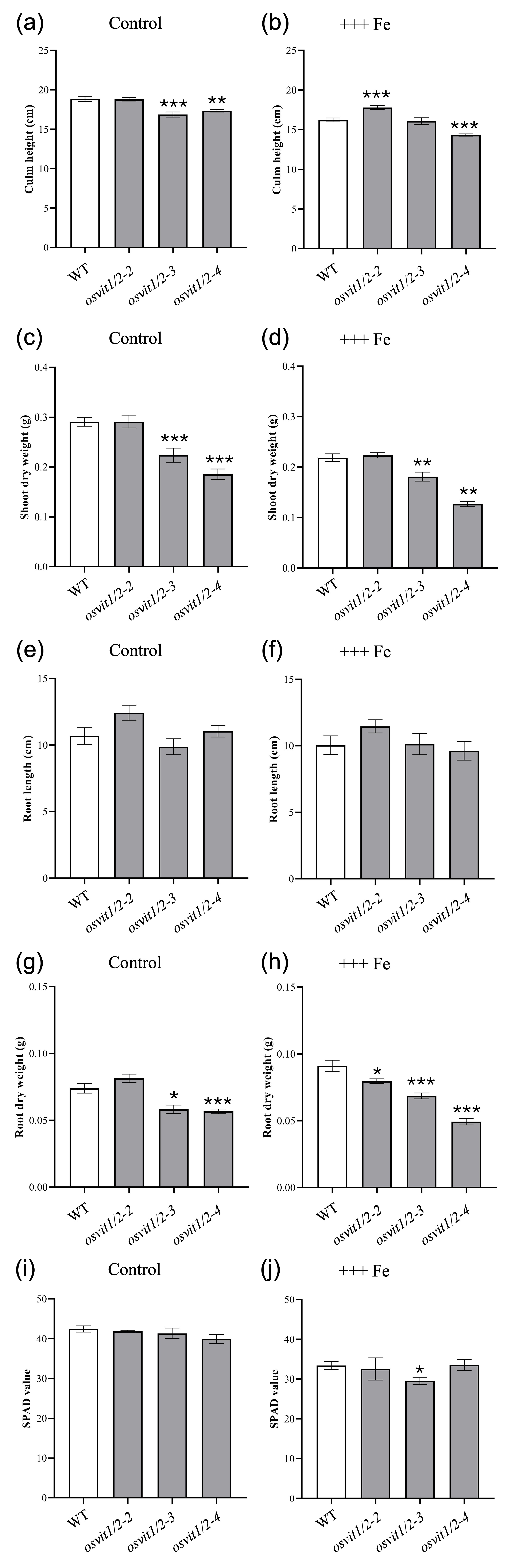

### Figure S4: Elemental profiling of WT and osvit1osvit2 leaves and roots of plants under control and iron excess.

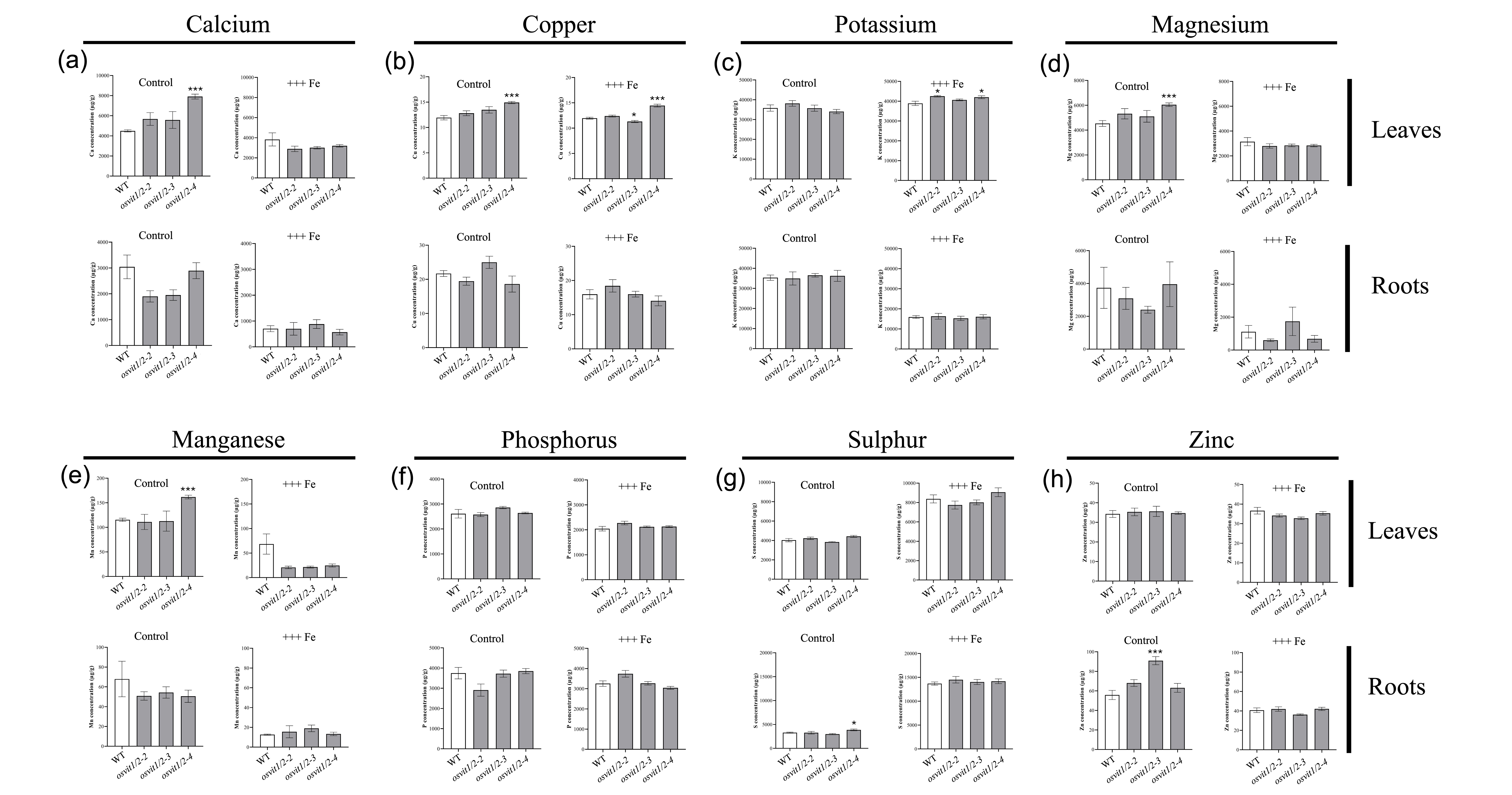

### Figure S5: Elemental profiling of WT and osvit1osvit2 seeds.

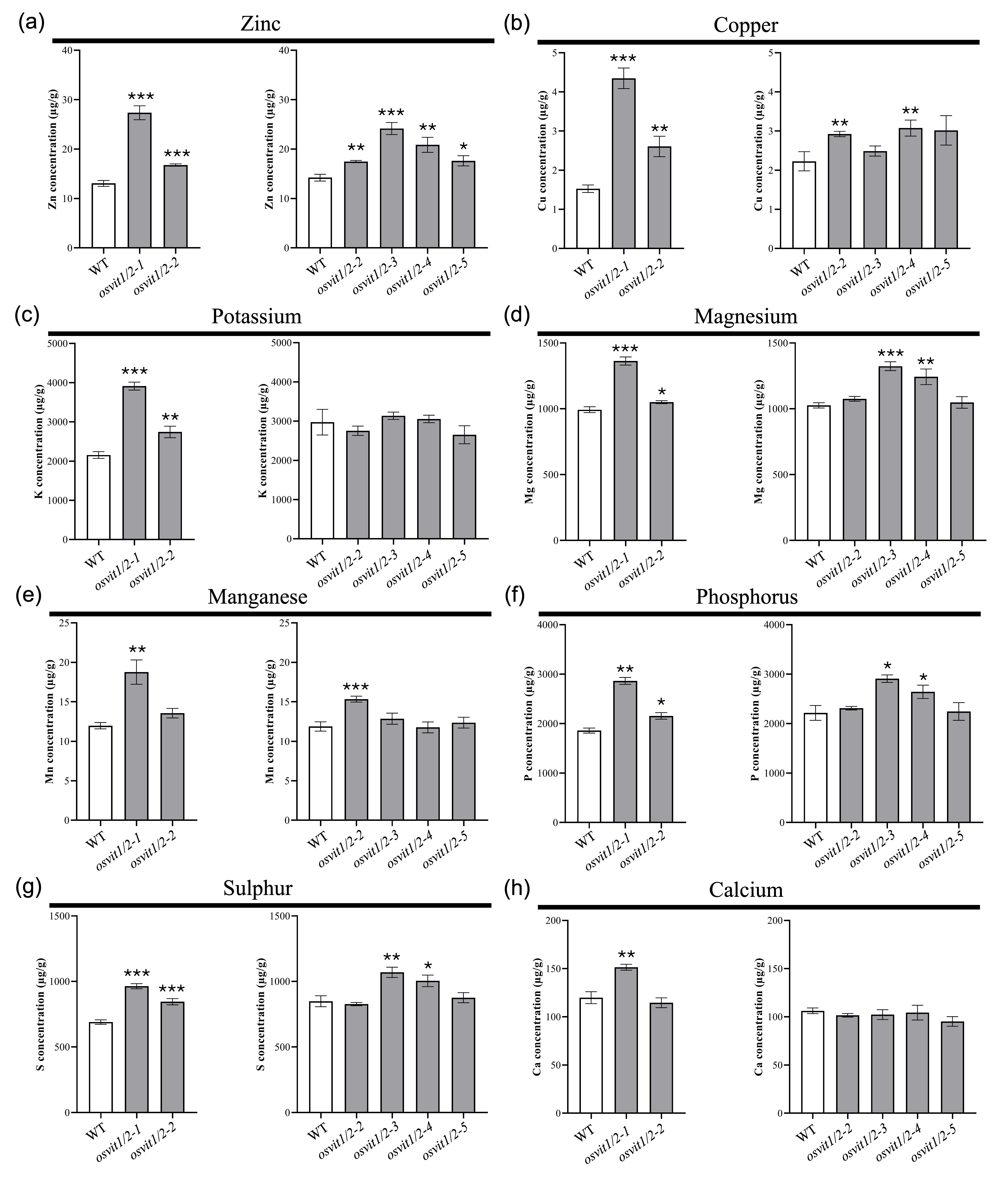

### Figure S6: osvit1osvit2 double knockout mutants changes in agronomic traits.

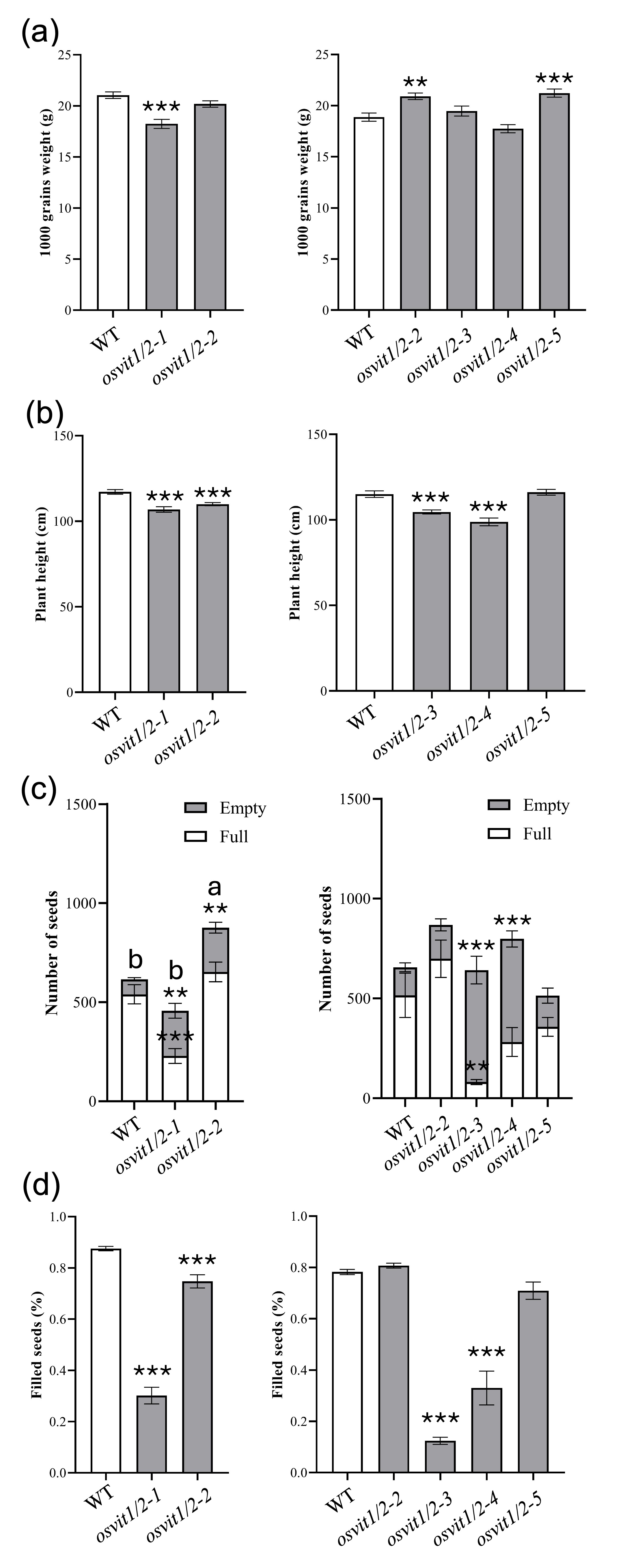

### Figure S7: Regions from WT and osvit1osvit2 embryos selected for high resolution analyses using Synchrotron X-Ray Fluorescence.

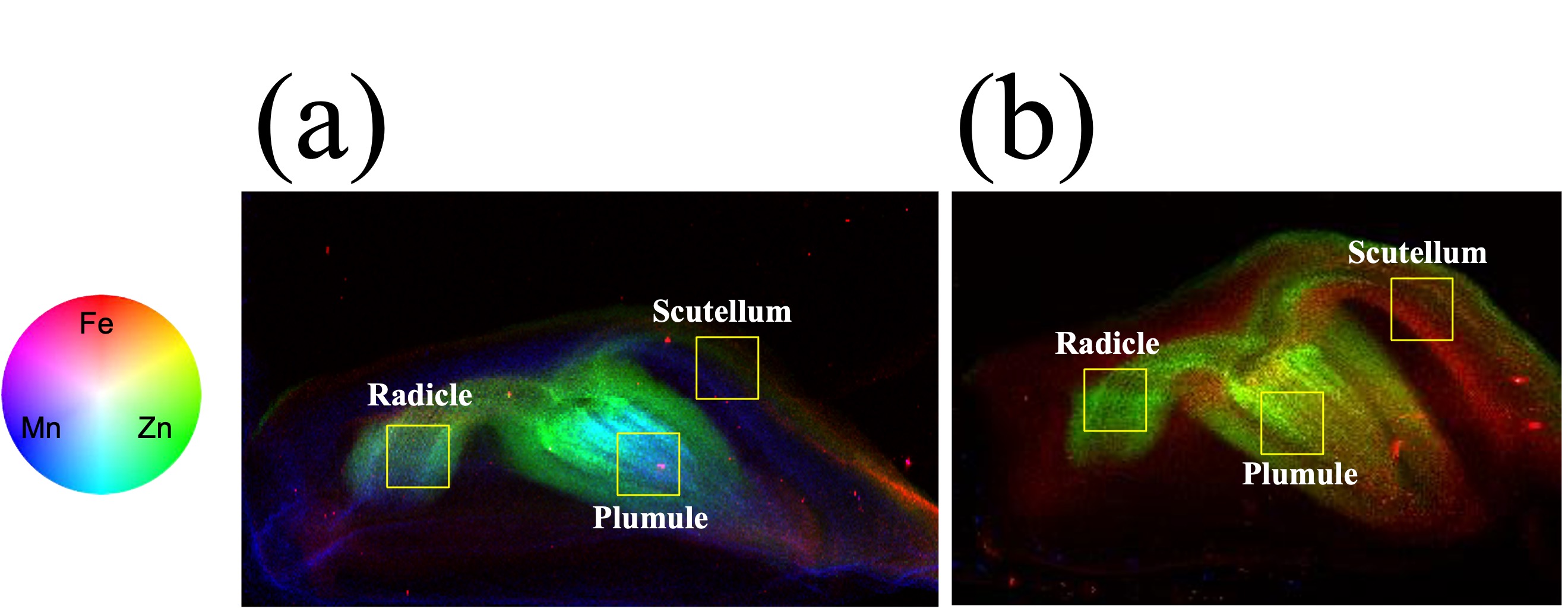

### Figure S8: Iron staining with Perls/DAB in aleurone layer cells.

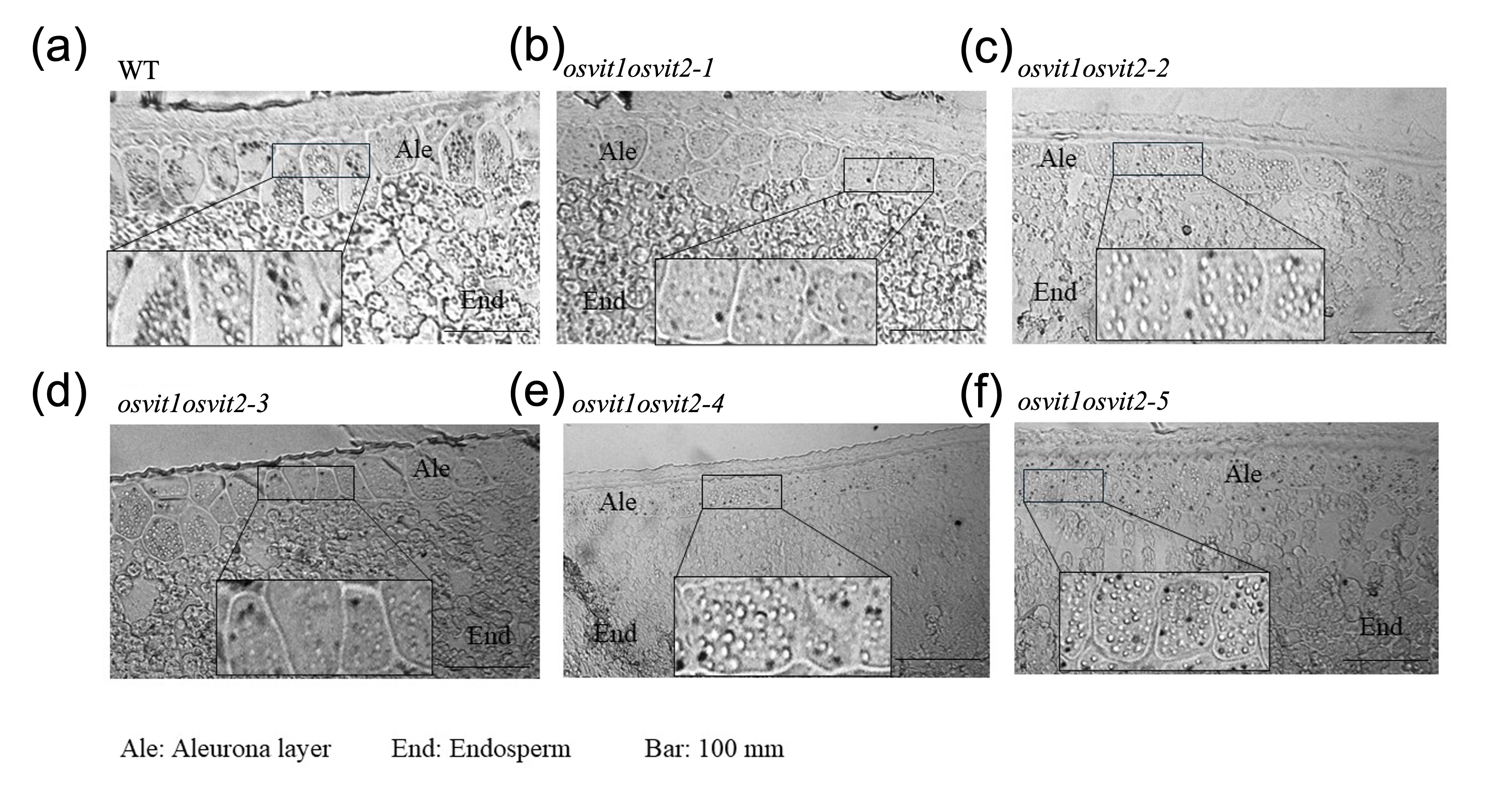
